# Quantitative Calibration of a Spatial QSP Model Identifies Fibroblast Impact on HCC Immunotherapy

**DOI:** 10.1101/2025.10.31.685939

**Authors:** Shuming Zhang, Hanwen Wang, Yeonju Cho, Wendy Wong, Mark Yarchoan, Elizabeth M. Jaffee, Won Jin Ho, Luciane T. Kagohara, Elana J. Fertig, Aleksander S. Popel, Atul Deshpande

## Abstract

Computational models are increasingly used to predict treatment response and optimize cancer therapy strategies. Among these, quantitative systems pharmacology (QSP) models mechanistically simulate tumor progression and pharmacological interventions, enabling virtual clinical trials, model-informed drug development, and biomarker identification. Coupling QSP with an agent-based model yields a spatial QSP (spQSP) platform that captures tissue-level spatial organization of the tumor microenvironment (TME). However, parameterizing such models to represent tumor biology remains an open problem. In this study, we developed a calibration framework using the Approximate Bayesian Computation - Sequential Monte Carlo (ABC-SMC) approach to calibrate the spQSP model with a combination of clinical and spatial molecular data, reflecting the TME characteristics of human tumors. This calibration framework matches tumor architectures between spQSP model predictions and patient spatial molecular data by fitting statistical summaries of cellular neighborhoods. We demonstrate that model calibration using CODEX data from untreated HCC patients enables prediction of TME spatial molecular states in an independent cohort receiving immune-checkpoint inhibitor (ICI) and tyrosine kinase inhibitor (TKI) combination therapy. Finally, we identify spatial and non-spatial pretreatment biomarkers and assess their predictive power for therapeutic response. This workflow demonstrates how integrating spatial-omics with multiscale mechanistic models enables quantitative calibration, biological insight, and in silico biomarker discovery, providing a framework for personalized cancer therapy across tumor types.

**Significance:** Digital twins and computational models are increasingly used to simulate disease and guide therapy, but they often struggle to capture the immense complexity driving the spatiotemporal evolution of the TME. The challenge is compounded by clinical sample limitations, which typically provide measurements from only static snapshots of the TME for parameter estimation. We demonstrate how mechanistic modeling frameworks can overcome this limitation by enabling inference of spatiotemporal model parameters from static spatial data – an intractable task for purely data-driven approaches. Ultimately, our work presents a workflow that integrates spatial-omics with multiscale mechanistic models, enabling quantitative calibration, deeper biological insight, and in silico biomarker discovery.

## Introduction

Computational models have emerged as powerful tools for predicting treatment response and optimizing therapeutic strategies. Mechanistic mathematical frameworks can encode known regulatory programs of tumor systems and be parameterized from clinical data to simulate tumor dynamics, immune interactions, and drug responses based on biological knowledge *in silico*. Of the family of mathematical models, quantitative systems pharmacology (QSP) models offer a mechanistic and multiscale representation of disease progression and pharmacological interventions ^1–6^, enabling virtual clinical trials, hypothesis testing, and model-informed drug development. By coupling an agent-based model (ABM) with a QSP model, we have built a spatial QSP (spQSP) platform to predict tumor dynamics at the organ level while reflecting spatial organization of the tumor microenvironment at the tissue level^7–10^, while making such predictions using clinical data alone is insufficient. However, as more mechanistic details are integrated into these computational models, systematic calibrations of model parameters are required for accurately predicting the spatiotemporal dynamics of the tumor microenvironment (TME).

Advances in spatial molecular characterization of the tumor microenvironment have occurred concurrently with advances in mechanistic mathematical models^11^. Spatial proteomic and transcriptomic data, with their high dimensionality and biological variability, provide rich information sources to parametrize mechanistic mathematical models of the tumor microenvironment^12^ and assess model fit^13^. However, these static snapshots of the TME pose a challenge when calibrating non-linear, stochastic models to capture the complex dynamics of the underlying biological processes in the TME. In our earlier work, we addressed this limitation by summarizing spatial information using spatial interaction metrics, allowing for qualitative model calibration based on spatial trends rather than exact cell locations^7,9^. Still, the qualitative nature of these approaches results in the need for manual tuning of parameters. Quantitative integration of spatial data with mechanistic models through data assimilation methods have been shown to increase spatial-temporal model accuracy in other scientific disciplines^14^, and are needed to enhance the accuracy of model-based predictions of complex systems-level response to cancer therapies.

In this study, we present a workflow for parameter inference of a mechanistic model of using spatial omics from clinical biospecimens and validation of model outputs. We apply this model to liver cancer, as it has a significant clincial impact projected to account for more than 42,000 new cancer cases and 30,000 cancer related deaths in the United States in 2025^15^, with hepatocellular carcinoma (HCC) comprising approximately 90% of all cases. Although systemic therapies have shown promising results with immunotherapies now making this disease treatable, only a small group of patients had long-term benefits^16^. With wide varity of intervention strategies, including tyrosine kinase inhibitors (TKI)^17,18^, immune-checkpoint inhibitors (ICI) ^19–21^, and cancer vaccines ^22^, matching patients with suitable therapies requires identifying predictive biomarkers ^23^. We introduce an expanded version of the spQSP model, which includes a fibroblast module to capture a key cell type we previously associated with resistance to immunotherapy in HCC ^21,24^. We then demonstrate how we quantiatively calibrated cell motility parameters in the spQSP model by matching predicted tumor architecture characteristics with those derived from treatment-naïve patient spatial proteomic data^25^. This calibration method was performed using the pyABC framework^26^, a Bayesian inference approach, with summary statistics of the cellular neighborhoods using Functional Cellular Neighborhood (FunCN)

scores^27^. The calibrated model is used to simulate the dynamic evolution of the TME under ICI and TKI combination therapy and the predicted tumor architectures are validated against post-treatment spatial proteomic data^21^ and spatial transcriptomics data^24^ from an HCC clinical trial^21^. Furthermore, we identify both spatial and non-spatial pretreatment biomarkers and quantify their predictive power for therapeutic response. Altogether, the workflow presented in this study can be applied to integrate spatial omics data into multiscale mechanistic models, enabling quantitative model calibration, biological interpretation, and in silico biomarker discovery for personalized cancer therapy beyond liver cancer.

## Results

### Approximate Bayesian Computation - Sequential Monte Carlo (ABC-SMC) Approach Provides a Framework for Parameter Inference in the spQSP Model

Our workflow has three primary components involving the construction, calibration, and validation of a model of liver cancer using the spQSP framework (Fig. 1A). The first component is an extended spQSP model incorporating a novel fibroblast module. Immunosuppressive cells such as M2-like macrophages, cancer stem cells, and regulatory T cells (Tregs) secrete transforming growth factor-beta (TGFβ), which promotes the activation of hepatic stellate cells (HSCs) into cancer-associated fibroblasts (CAFs). The second, a model calibration component, uses an Approximate Bayesian Computation - Sequential Monte Carlo (ABC-SMC) approach to calibrate QSP parameters and ABM parameters in spQSP model separately. Specifically, we fitted QSP parameters using experimental data and fitted ABM parameters using CODEX from Qiu et al ^25^. Finally, the model validation component compares our model outputs with spatial omics from post-treatment clinical biospecimens.

**Fig. 1.**
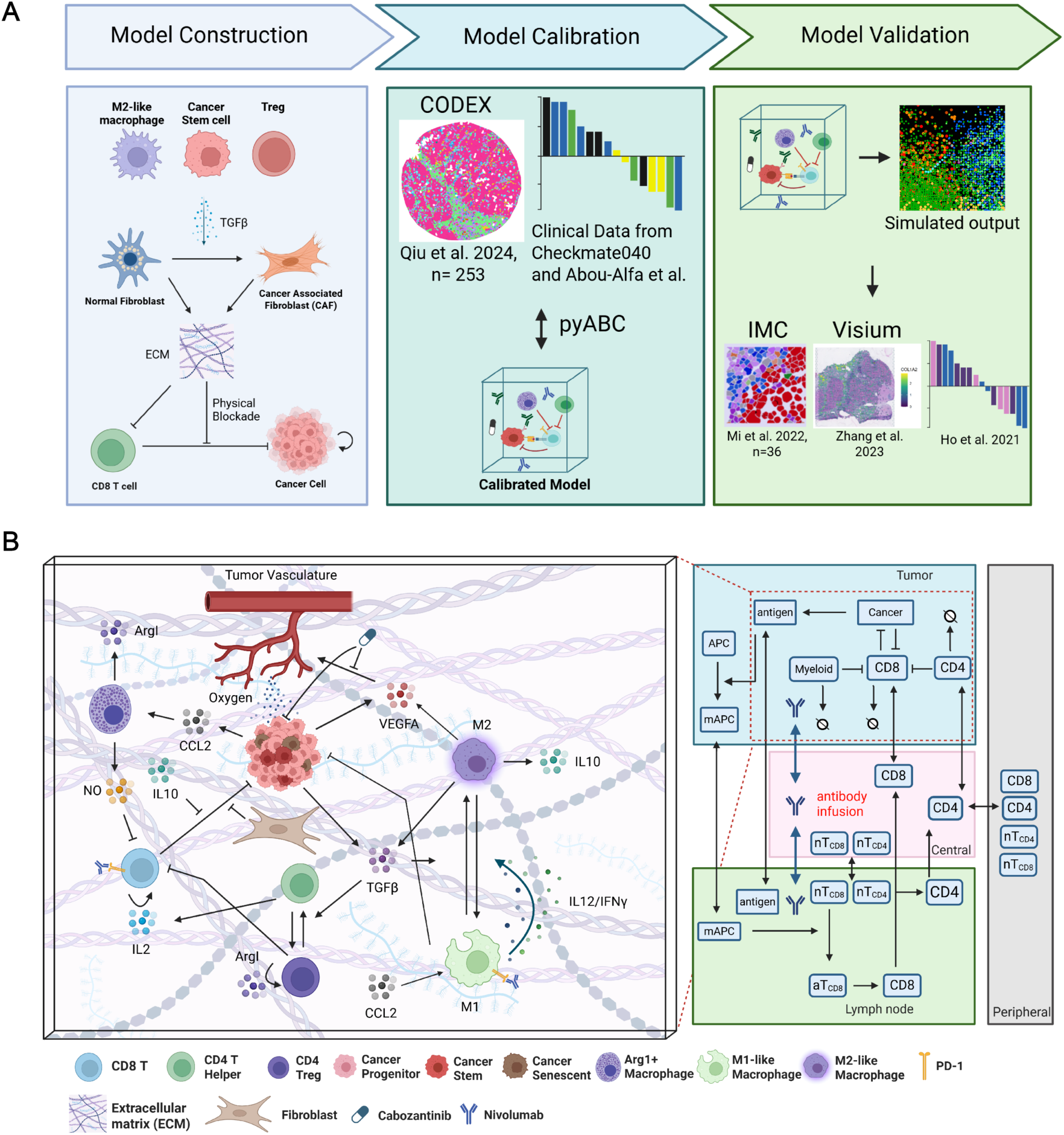
Overview and Model Schematics. **(A)** Overview of model construction, calibration using pre-treatment CODEX data and clinical trials, and validation with post-treatment IMC, Visium data, and outcomes. **(B)** Spatial QSP model combining QSP and ABM to simulate immune-tumor interactions and cytokine diffusion in the tumor microenvironment.

The spatial quantitative systems pharmacology (spQSP) model is built upon the robust framework of our previously developed for hepatocellular carcinoma (HCC)^7^. The spQSP model is designed to simulate clinical data at both the organ and tissue scales and consists of two coupled submodels: a non-spatial, whole-body QSP model and a spatially resolved agent-based model (ABM). The QSP component is formulated as a system of ordinary differential equations (ODEs) that captures cellular dynamics, trafficking, anti-tumor immune responses, as well as the pharmacokinetics (PK) and pharmacodynamics (PD) of targeted therapies and immunotherapies (Supplemental Fig. 1). The ABM component captures the spatial architecture of the tumor microenvironment (TME) (Fig. 1B). It comprises two layers: a cellular layer that tracks the type, state, and spatial location of each individual cell; and a molecular layer that resolves local concentrations of diffusible factors (e.g., cytokines) using partial differential equations (PDEs).

A challenge of the multi-scale modeling framework is to calibrate the parameters of these models to fit the human TME and therapeutic response. In previous work, we have based this calibration exclusively on clinical data and demonstrated qualitive fits of the spatial ABM against spatial molecular data. In this study, we will calibrate QSP model parameter first against clinical data and subsequently develop new methods to further calibrate the ABM directly with spatial molecular data while retaining the coupling with the QSP model.

### Model Calibration, Virtual Clinical Trial, Biomarker Identification in QSP Model

In this section, we calibrate the non-spatial parameters in the QSP model using ABC-SMC approach implemented in the pyABC framework^26^ (Fig 2A). The calibrated model simulates a virtual clinical trial and identifies potential biomarkers of immunotherapy responses. After constructing the cabozantinib module and data assimilation framework, we calibrate cancer cell growth rates kc,growth, vascular expansion rates kK,g, and cabozantinib inhibition rates kK,cabo using tumor volume dynamics from experimental mouse data using the pyABC framework26,28. Specifically, k and kK,cabo were estimated using control group data (Supplemental Fig. 2A-F), while kK,cabo was fitted using data from the cabozantinib treatment group until convergence (Supplemental Fig. 2G-J). Since the model parameters were initially calibrated using mouse data, they were subsequently converted to human-equivalent values through allometric scaling, using a scaling factor of, based on the mass ratio between human and mouse^29^. The calibrated parameters from the murine data were then translated to human-equivalent values through allometric scaling. Existing QSP parameters were adapted from our previously published HCC model^6^.

**Fig. 2.**
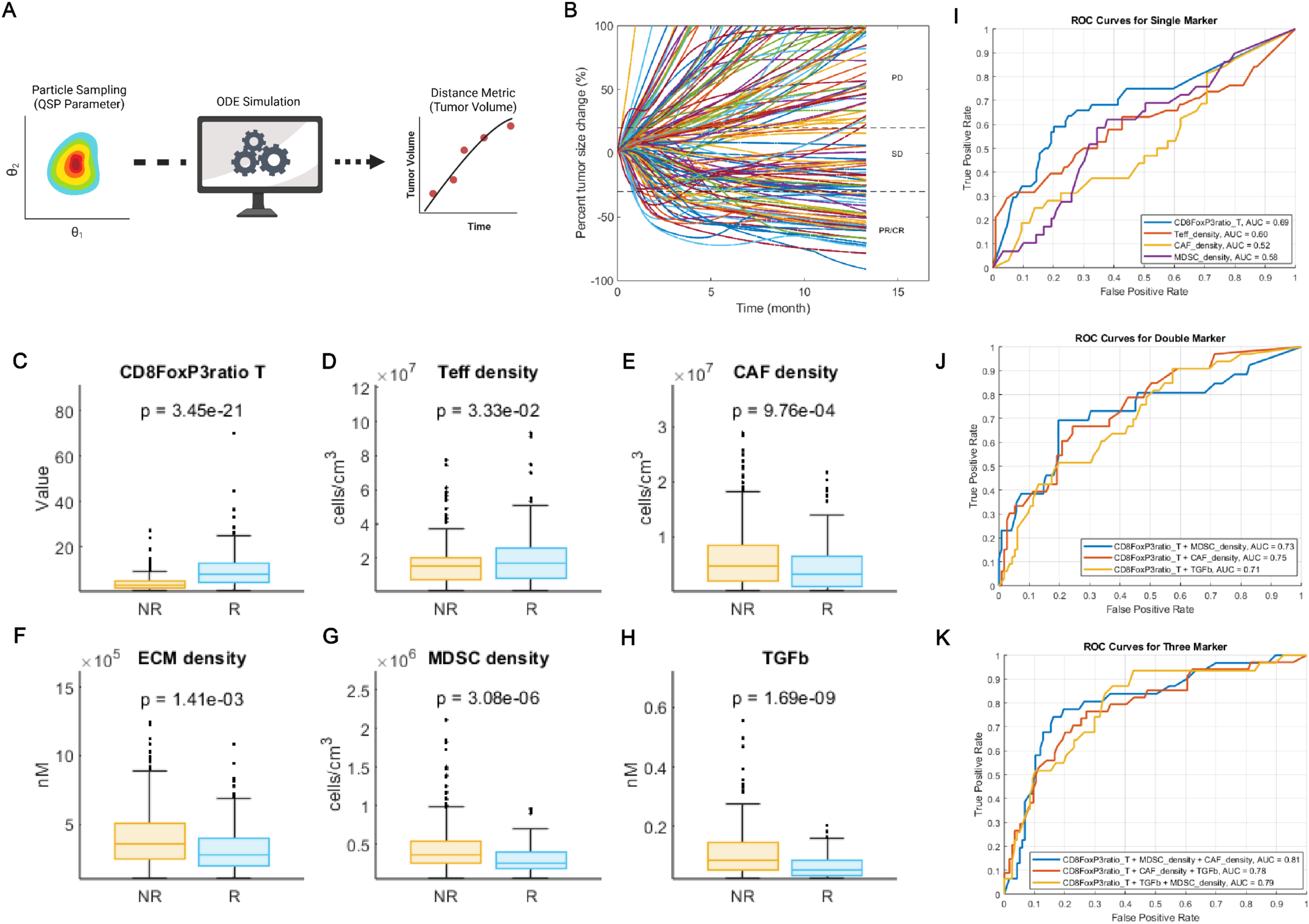
QSP calibration and simulation results. (A) Workflow of QSP parameter inference using ABM-SMC approach implemented in pyABC. **(B)** Spider plot showing tumor size change over time for 150 randomly selected virtual patients out of 500. **(C–H)** Differences in pre-treatment biomarkers between virtual responders (R) and non-responders (NR): (C) CD8/FoxP3 ratio in the tumor, (D) Teff cell density, (E) CAF density, (F) ECM density, (G) MDSC density, and (H) TGFβ concentration. P-values are shown for each comparison. **(I–K)** Receiver Operating Characteristic (ROC) curves evaluating the predictive power of: (I) single markers, (J) pairwise marker combinations, and (K) triple marker combinations from panels (C–H). Area under the curve (AUC) values are indicated, 70% of the virtual cohort used for training, and 30% used for testing.

To validate the calibrated model, we simulated cabozantinib monotherapy in a virtual cohort of 500 patients. In accordance with the clinical trial protocol, we assessed patient response using the RECIST 1.1 criteria. Our virtual cohort yielded an objective response rate (ORR) of 3% (95% CI: 0–6%; Supplemental Fig. 3A, B), consistent with results from a phase III clinical trial ^30^. Subsequently, Nivolumab-related pharmacodynamic parameters (see Supplemental Table) were also adopted from previous model^6^. The ORR from simulated nivolumab monotherapy matches the ORR observed in the CheckMate 459 trial^19^ (ORR 17.1% in simulation vs. 15% in CheckMate 459, Supplemental Fig. 3C, D). For model validation, we simulated combination therapy with cabozantinib and nivolumab in the same virtual patient cohort (n = 500). The simulation outputs were subsequently validated using ORR data from the nivolumab plus cabozantinib combination therapy cohort (Cohort 6) in the CheckMate 040 trial^20^. The predicted ORR was 21.6% (95% CI: 13.0–30.0%), closely matching the 18.1% ORR reported in CheckMate 040 Cohort 6 ^20^ (Fig. 2B, Supplemental 3E). We further validated the model by analyzing tumor microenvironment features between responders and non-responders (Fig. 2C–H). The simulated T cell density was consistent with clinically reported values ^21^. Responders exhibited higher Teff (CD8 T cell) density and Teff-to-Treg ratio compared to non-responders (Fig. 2C, D), consistent with findings from CheckMate 040^31^. Similarly, fibroblast and extracellular matrix (ECM) densities were elevated in non-responders (Fig. 2E, F), in agreement with clinical observations^32^ (Fig. 2E, F). MDSC density and TGFβ concentration were also higher in non-responders, suggesting their immunosuppressive role within the tumor microenvironment^33^ (Fig. 2G, H). To further evaluate the model’s ability to predict patient outcomes, we trained logistic regression classifiers using combinations of spatial biomarkers. Among single biomarkers, CD8/FoxP3 ratio showed the highest predictive power (AUC = 0.69), outperforming Teff, MDSC, and CAF densities (AUCs = 0.60, 0.58, and 0.52, respectively).

Combining markers improved classification performance: the best-performing double-marker model—CD8/FoxP3 ratio with CAF density—achieved an AUC of 0.75. Triple-marker models yielded the highest predictive accuracy, with the top combination (CD8/FoxP3 ratio + MDSC + CAF densities) reaching an AUC of 0.81. These results demonstrate the model’s potential to prospectively identify predictive biomarkers for immunotherapy response prior to clinical trials.

### Neighborhood Analysis Identifies Association between Fibroblast-Dense Tumor Architectures and Poor Overall Survival

The calibrated QSP model contains only non-spatial attributes and does not reflect the spatial architecture of the TME. Prior to expanding this approach to a calibrated spatially resolved model, we first sought to quantify the cellular architecture of the TME from spatial molecular data of human HCC as a baseline prior to in silico modeling. Briefly, Functional Cell Neighborhood (FunCN) is a novel spatial metric that estimates cell-cell influence through a spatial kernel-based approach^27^. This metric was selected for our study because FunCN score can capture cell influence over the entire region of interest measured with spatial molecular data.

Prior to model calibration, we first analyzed publicly available CODEX imaging data from Qiu et al.^25^ to determine the clinically relevant featured needed for accurate model calibration and to ensure the robustness of this dataset for spatial model calibration. This dataset includes 401 tissue microarray (TMA) samples from 401 treatment-naïve hepatocellular carcinoma (HCC) patients. To reduce tissue selection bias, we included only samples with a cancer cell fraction greater than 0.5 for downstream analysis; cell segmentation results for a short-term survivor (Fig. 3A, *TMA3_8_reg013*) and a long-term survivor (Fig. 3B, *TMA3_8_reg032*) are shown. For each selected sample, we computed the FunCN scores (Fig. 3C; see Methods). Corresponding FunCN score matrices of the example short-term and long-term survivors (Fig. 3D, E) are shown. We identified five FunCN scores significantly associated with long term patient survival among cell types implemented in our spQSP model. We define FunCN(Target, Source) as the average spatial influence of source cell type on a target cell type based on their local spatial proximity. Among these, FunCN(CD8, Tumor) and FunCN(CD4, Tumor) were positively correlated with overall survival, marking them as favorable prognostic indicators (Fig. 3F, G). Conversely, FunCN(Fibroblast, Fibroblast), FunCN(Treg, Treg), and FunCN(CD8, Macrophage) were negatively associated with survival (Fig. 3H–J, Supplemental Fig. 4), indicating immunosuppressive or stromal exclusion phenotypes. We further consolidate our findings by examining overall survival divided by treatments (Supplemental Fig. 5). Notably, patient sample *TMA3_8_reg013* exhibited a high FunCN(Fibroblast, Fibroblast) score and low FunCN(CD8, Fibroblast) score and was classified as a short-term survivor (overall survival = 59 days). In contrast, sample *TMA3_8_reg032*, with low FunCN(Fibroblast, Fibroblast), representing low clustering, and high FunCN(CD8, Tumor), corresponded to a long-term survivor (overall survival = 2089 days). These five FunCN scores were subsequently compiled across all samples to inform and calibrate the spatial modules of the spQSP model (Fig. 3K, Supplemental Fig. 6). These observations expand our association of CAFs with poor immune function in our smaller spatial transcriptomics cohort^24^, further justifying the extension of our previous mathematical model^7^ to include CAFs and the calibration of their relative spatial neighborhoods in this current study.

**Fig 3.**
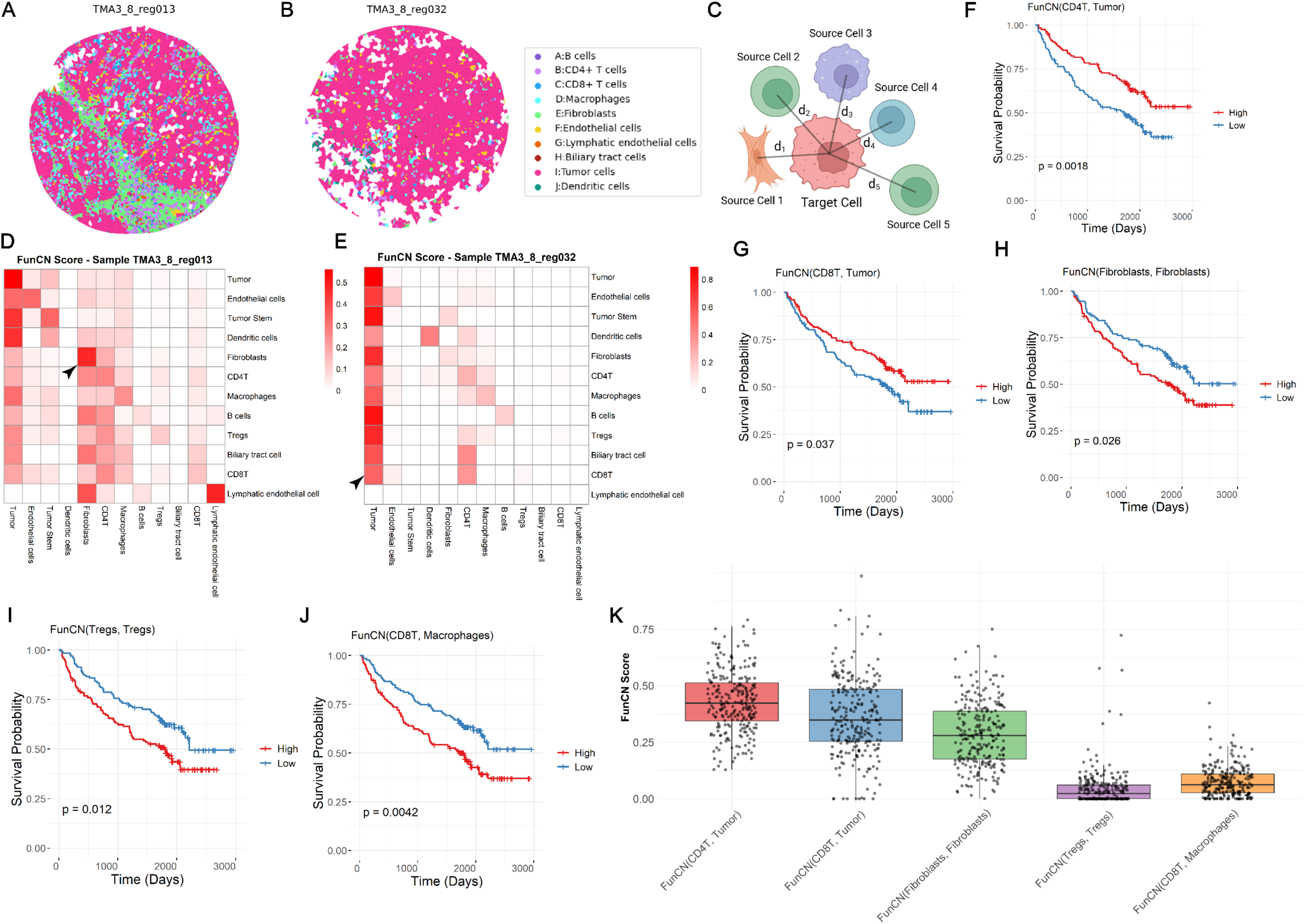
FunCN analysis on Pre-treatment CODEX data. **(A, B)** Example cell phenotyping maps from Qiu et al. showing a short-term survivor (A) and a long-term survivor (B). Cell types are color-coded as indicated. **(C)** Schematic illustrating the calculation of the FunCN score for a target cell based on its surrounding source cells and their spatial distances. **(D, E)** FunCN score matrices for the samples shown in panels A and B, respectively. Rows represent target cell types, and columns represent source cell types. The color intensity corresponds to the magnitude of the FunCN score. **(F–J)** Kaplan-Meier survival curves for five selected FunCN metrics, including FunCN(CD8, Tumor), FunCN(CD4, Tumor), FunCN(Fibroblast, Fibroblast), FunCN(Treg, Treg), and FunCN(CD8, Macrophage) that significantly stratify patient overall survival. Patients are grouped into high vs. low FunCN score cohorts, with adjusted p-values calculated using the False Discovery Rate (FDR). **(K)** Boxplots summarizing the distributions and medians of the five selected FunCN scores used for spQSP model calibration. Each point represents a patient sample.

### ABM Parameter Inference Based on Pre-treatment Tumor Architecture

After the calibration and validation of the QSP model based on clinical data, we coupled the ABM with calibrated QSP model, forming a new spQSP model (Fig 1B). Then spatial parameters in the spQSP model were calibrated based on the similarity in tumor architecture between simulated tumors and the CODEX imaging data from patients quantified by FunCN scores. Spatial parameters related to cellular motility that are critical to model function cannot be directly measured just from measurements of static tumor microenvironment states, as captured by CODEX. Nonetheless, we demonstrate the ability to calibrate these parameters by leveraging the pyABC framework and using FunCN scores as a common statistical summary metric to assess the similarity of model-predicted tumor architectures with those observed in the CODEX data from human biospecimens. Given their association with patient outcomes, the five FunCN scores used for model calibration include spatial, pairwise relationships between cancer, T cell, fibroblasts, and macrophages, which contain all cell modules implemented in the spQSP model. To compare the results with two-dimensional CODEX data, we slice the ABM output along y-axis, calculate the FunCN score, and take the average of the score to represent the simulated tumor architecture. We calibrated cell motility parameters in the ABM, including those for T cells, fibroblasts, and macrophages, through an ABC-SMC approach^26^ in pyABC (Fig 4A) to match the tumor architectures from the CODEX data based on selected spatial metrics (Fig 3K).

**Fig. 4.**
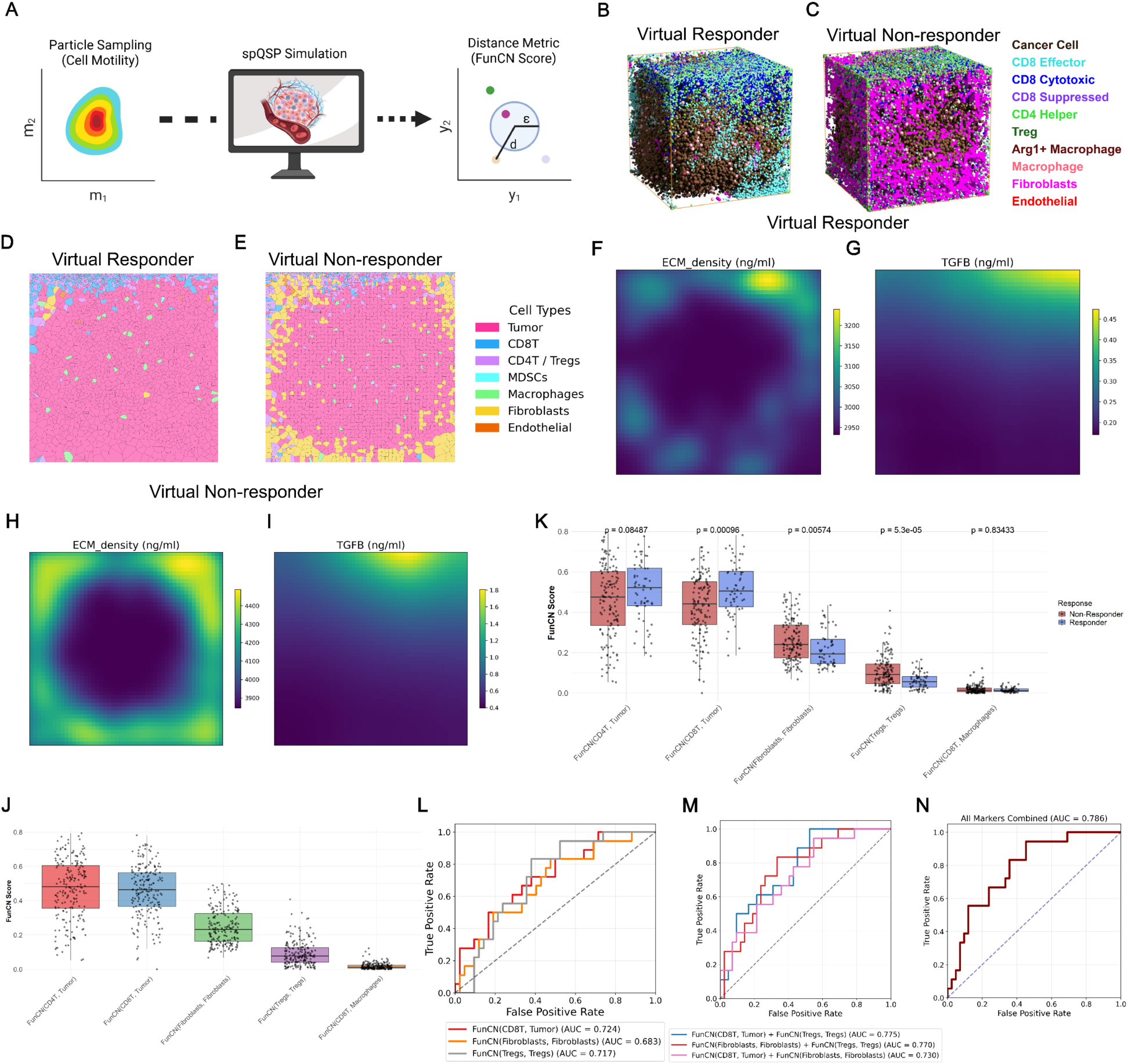
| Pre-treatment (Day 0) spatial analysis on simulated tumor architecture after model calibration. **(A)** Schematics of calibrating cell motilities using pyABC platform **(B, C)** Three-dimensional agent-based model (ABM) outputs from the spQSP model showing a virtual responder (B) and a virtual non-responder (C). **(D, E)** Voronoi plots showing spatial cell distributions for the same virtual responder and non-responder shown in panels A and B. Slices are taken along the y-axis at y = 25 for direct comparison with CODEX data. **(F–I)** Spatial heatmaps of extracellular matrix (ECM) density and TGFβ concentration for the virtual responder (F, G) and non-responder (H, I), corresponding to the same spatial slice as in panels D and E (y = 25). **(J)** Distributions of five FunCN scores across the calibrated virtual patient cohort shown as boxplots. **(K)** Boxplots showing the same five FunCN scores stratified by treatment response (Responder vs. Non-responder) at the pre-treatment stage. P-values are shown for each comparison. **(L-N)** Receiver Operating Characteristic (ROC) curves evaluating the predictive performance of FunCN metrics in stratifying response: (L) single FunCN markers, (M) pairwise marker combinations, (N) triple marker combinations. Area under the curve (AUC) values are shown. A 70:30 train-test split was used for model evaluation.

After calibrating the ABM parameters (Supplemental Fig. 7), we generated 200 virtual patients by sampling from the posterior parameter distributions of the calibrated spQSP model. These virtual patients were simulated under a combination immunotherapy regimen of nivolumab (240 mg, intravenously every two weeks) and cabozantinib (40 mg, orally daily) for 70 days, consistent with the dosing schedule in a reported phase 1b clinical trial^21^. Using the same clinical response criteria^21^, the simulated response rate was 30.3% (30/198), excluding 2 virtual patients due to implausible parameter sets. This response rate aligns closely with the clinical trial outcome of 35.7%. To assess the fidelity of the calibrated model in reproducing spatial tumor-immune architecture, we visualized three-dimensional cell distributions for a virtual responder and non-responder at pre-treatment (Fig. 4B, C).

For direct comparison with CODEX, which provides two-dimensional spatial profiles, we also generated 2D cell distributions for the same virtual patients (Fig. 4D, E). Notably, the virtual non-responder exhibited more pronounced fibroblast clustering around tumor cells, along with extracellular matrix (ECM) density and TGFβ concentration (Fig. 4F–I), consistent with the known association between CAFs and poor prognosis^34^. Furthermore, population-level comparisons of FunCN scores, quantifying spatial colocalization between cell types, revealed consistency between simulations and CODEX data (Fig. 3K, 4J, Supplemental Fig. 8). Our model recapitulated key spatial features, identifying FunCN(CD8, Tumor) and FunCN(CD4, Tumor) as positive prognostic markers, while FunCN(Fibroblast, Fibroblast) and FunCN(Treg, Treg) were associated with poor prognosis, under the assumption that patients with longer overall survival respond better to systemic therapy from Qiu et al.^25^ (Fig. 4K). Finally, to evaluate the predictive power of these five FunCN scores, we constructed logistic regression models using each score individually and in combination. The best single-score model achieved an AUC of 0.724, while the top-performing two-score combination reached an AUC of 0.775. Incorporating all five markers resulted in a maximal AUC of 0.786 (Fig. 4L–N), demonstrating the predictive value of spatial features in stratifying therapeutic response.

### Integrative Validation of spQSP Model with IMC and Visium Data Highlights Post-Treatment Spatial Remodeling

To further validate the spQSP platform, we compared post-treatment simulation results with independent spatial molecular data from patients treated with cabozantanib and nivolumab, including Imaging Mass Cytometry (IMC) proteomic data^21,33^ and Visium spatial transcriptomics^24^. In the simulations, virtual responders exhibited physical contact between infiltrating lymphocytes and tumor cells, characterized by FunCN(CD8, Tumor) and FunCN(CD4, Tumor), key features of effective anti-tumor immunity. In contrast, virtual non-responders displayed fibroblast-driven spatial segregation, wherein fibroblasts acted as physical barriers that separated T cells from cancer cells, thereby limiting immunotherapy efficacy (Fig. 5A–D). In addition, ECM density and TGFβ concentration are shown with spatial resolution for virtual responders and virtual non-responders (Fig. 5E–H). Consistent with pre-treatment observations, post-treatment samples from non-responders demonstrated elevated ECM density and TGFβ concentration (Fig. 5I, J), highlighting the persistent role of stromal remodeling in therapeutic resistance.

**Fig. 5.**
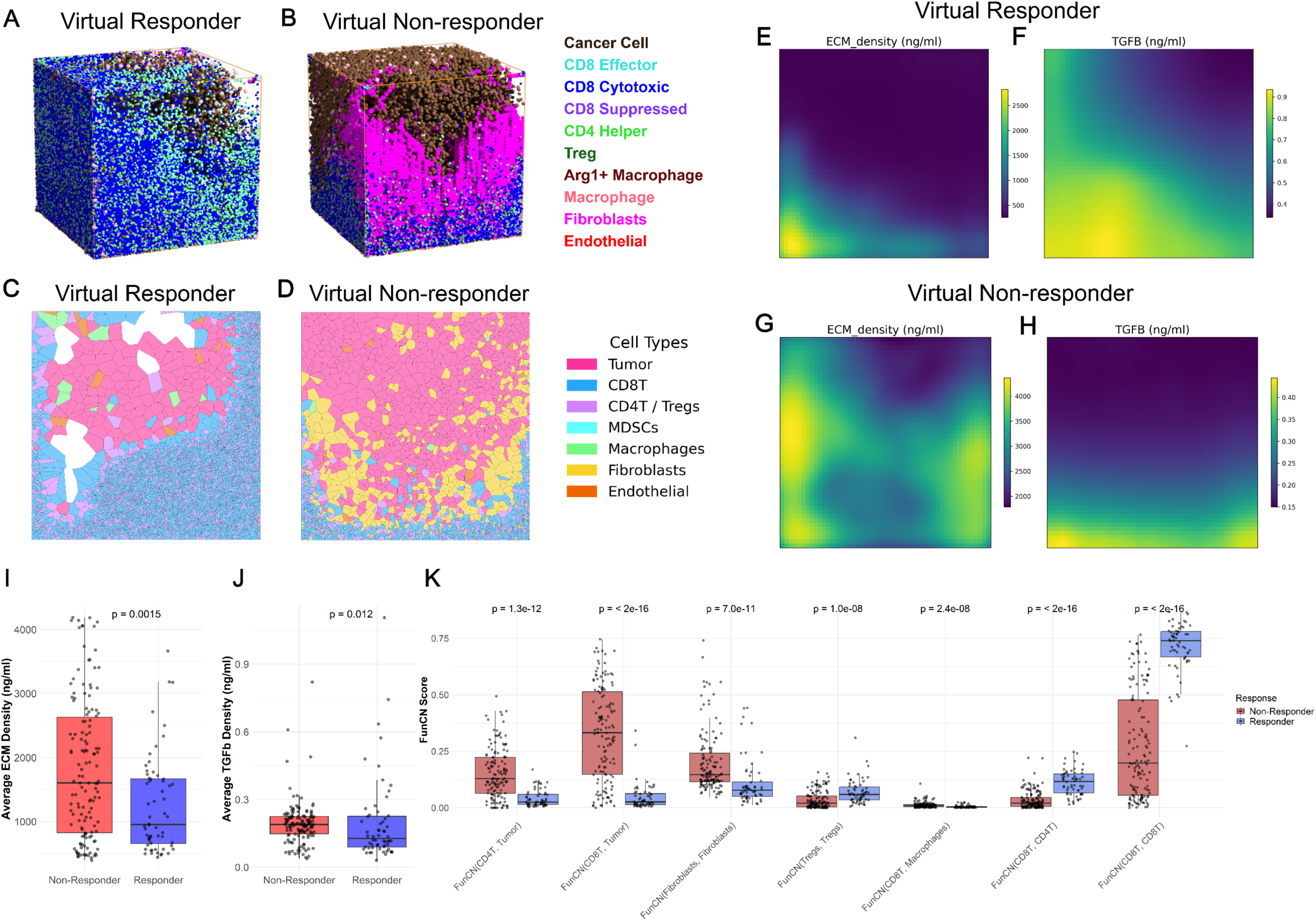
Post-treatment (Day 70) spatial analysis on simulated tumor architecture for model validation. **(A, B)** Three-dimensional ABM outputs from the spQSP model showing a virtual responder (A) and virtual non-responder (B). These simulations correspond to the same virtual patients shown in Fig. 4A and 4B. **(C, D)** Voronoi diagrams representing cell distributions for the virtual responder and non-responder from panels A and B, respectively. Slices are taken along the y-axis (y = 25). **(E–H)** Spatial distribution maps of ECM density and TGFβ concentration for the virtual responder (E, F) and non-responder (G, H), corresponding to the same spatial slice shown in panels C and D. (I, J) Average ECM and TGFβ density for virtual responder and virtual non-responder at post-treatment, respectively. **(K)** Boxplots of seven post-treatment FunCN scores stratified by treatment response (Responder vs. Non-responder). P-values indicate statistical significance for each metric.

At the population level, the model also predicted an inverse trend in the post-treatment tumor architectures, as quantified by FunCN, compared to those seen in pre-treatment. Specifically, the predicted post-treatment FunCN(CD8, Tumor) and FunCN(CD4, Tumor) scores were reduced in virtual responders relative to non-responders (Fig. 5J), a result attributed to effective tumor clearance in responders, which decreases the opportunity for immune-tumor interactions.

Notably, post-treatment Treg aggregation was higher in responders than in non-responders— again reversing the pre-treatment pattern. Meanwhile, CAF clustering remained consistently elevated in non-responders at both timepoints, underscoring the immunosuppressive role of CAFs (Fig. 5J, Supplemental Fig. 9, 10).

These predicted post-treatment tumor architectures from the simulation results are corroborated by post-treatment tumor architectures in the IMC data. Clinical non-responders (Fig. 6A) exhibited lower lymphocyte infiltration compared to responders (Fig. 6B), closely recapitulating the spatial patterns predicted by the spQSP model. Application of FunCN to the IMC dataset revealed significantly lower FunCN(CD8, Tumor), FunCN(CD4, Tumor), and FunCN(Treg, Treg) scores in responders (Fig. 6C–E), in agreement with model predictions. These reductions are likely attributable to effective tumor clearance, which diminishes both the spatial proximity and interaction frequency between immune cells and tumor cells, resulting in decreased FunCN scores. Furthermore, two metrics that were not predictive at the pretreatment stage in the CODEX dataset—FunCN(CD8, CD4) and FunCN(CD8, CD8)—were significantly elevated in responders post-treatment. This suggests that enhanced lymphocyte–lymphocyte interactions, particularly among CD8+ and CD4+ T cells, are associated with favorable outcomes (Fig 5J, Fig. 6F, G). These findings align with emerging evidence that lymphocytes aggregation plays a critical role in mediating effective responses to immunotherapy^35,36^.

**Fig. 6.**
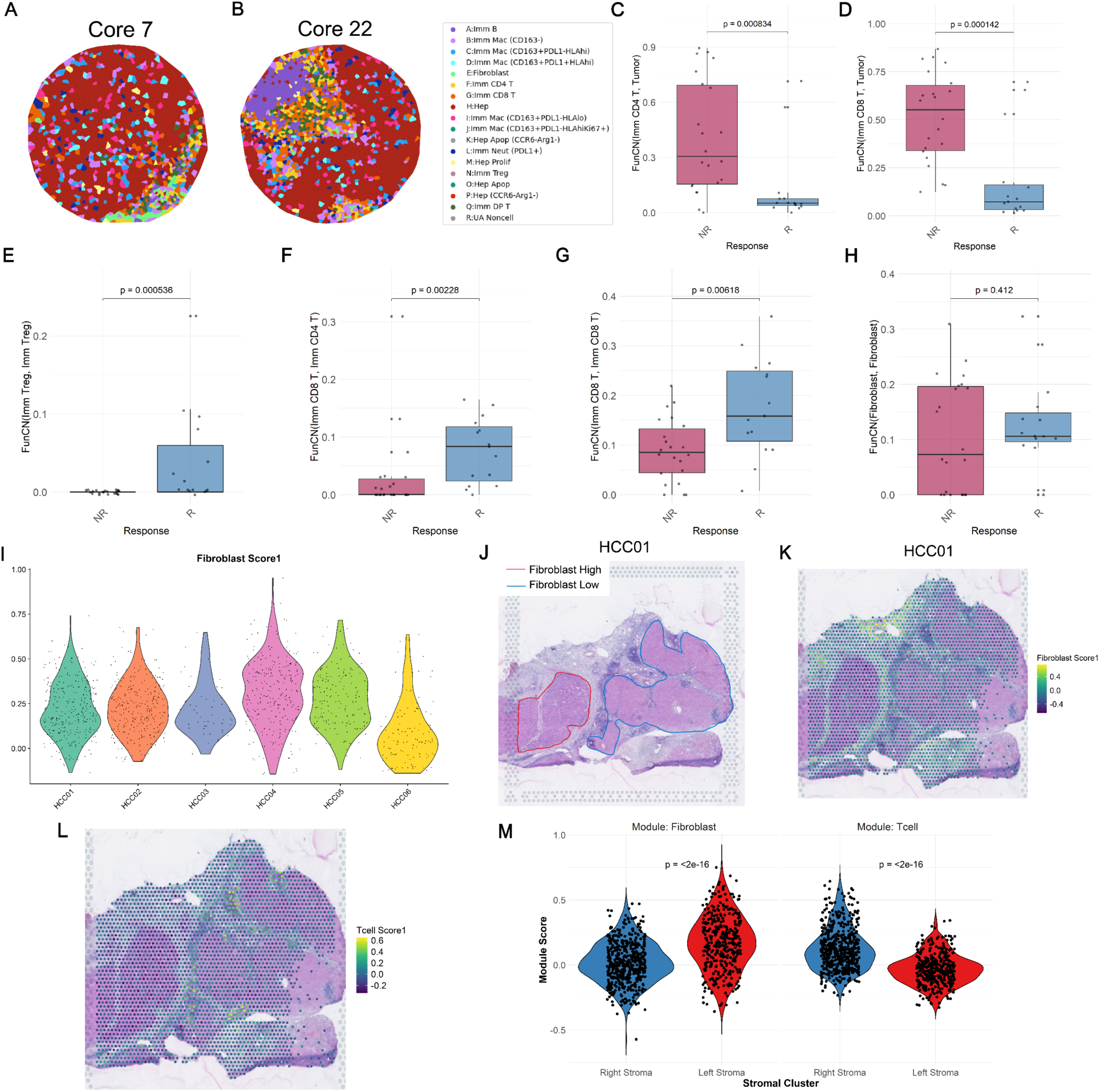
Post-treatment clinical sample spatial analysis on tumor architecture for model validation. **(A, B)** Representative Imaging Mass Cytometry (IMC) data from tissue microarrays (TMAs) for (A) a non-responder and (B) a responder, both treated with nivolumab plus cabozantinib combination therapy. Cell phenotypes are indicated in the legend. **(C–H)** Boxplots comparing post-treatment FunCN scores between responders (R) and non-responders (NR). P-values are shown above each panel. **(I)** Violin plots showing panCAFScore1 distributions across six Visium samples. Responders: HCC01–HCC04; Non-responders: HCC05–HCC06. **(J)** H&E-stained images from Visium slides showing manually outlined tumor regions with high (red) and low (blue) fibroblasts in Sample HCC01. **(K, L)** Spatial distribution of CAF-related module scores (panCAF Score1) and T cell module scores (T cell Score1), overlaid on the corresponding tissue sections in Panel J. CAF module genes: *LUM, DCN, COL1A1, VIM, ENTPD1, ACTA2, PDPN, COL1A2, SERPINF2*; T cell module genes: *CD3D, CD3E, CD8A, CD4, PTPRC, SELL, IL7R, CCL19, CCL21, GZMB.* **(M)** Violin plots showing CAF markers and T cell marker differences in fibroblast high and fibroblast low stroma.

In contrast to our simulation results (Fig. 5J), FunCN(Fibroblast, Fibroblast) scores did not significantly differ between responders and non-responders in the IMC dataset (Fig. 6H). Visium spatial transcriptomics data showed similar results (Fig. 6I; Responders: HCC01–04; Non-responders: HCC05, HCC06). However, upon examining individual post-treatment samples (e.g., HCC01), we identified two distinct tumor regions: one characterized by fibroblast dense region and the other by fibroblast light region (Fig. 6J, K). The fibroblast dense region has poor T cell infiltration compared to the fibroblast light region (Fig. 6L, M). Notably, the patient HCC01 responded to immunotherapy but experienced rapid relapse. A similar pattern was observed in an independent non-responder sample from Liu et al.^37^, where dense fibroblast regions also corresponded to limited immune cell infiltration (Supplemental Fig. 11). Importantly, in both samples, fibroblast-dense regions were located adjacent to tumor regions, consistent with spatial characteristics predicted by our simulation (Fig. 5D). These findings suggest that CAFs may physically restrict T cell access to tumor cells, thereby limiting contact-dependent immune responses and contributing to resistance to immunotherapy. To summarize, these results support both the predictive validity and mechanistic plausibility of the spQSP platform in recapitulating spatially resolved features of immunotherapy response.

### Investigate Relationship between Model Parameters and FunCN scores using Partial Rank Correlation Coefficient (PRCC)

To interrogate the relationships between model parameters and both spatial and non-spatial response metrics in virtual patients, we performed Partial Rank Correlation Coefficient (PRCC) analysis. PRCC was conducted separately for pre-treatment and post-treatment metrics (Fig. 7A, B). At the pre-treatment stage, the activation rate of cancer-associated fibroblasts (*k_caf_tran*) showed a positive correlation with FunCN(Fibroblast, Fibroblast) and CAF density, and a negative correlation with FunCN(CD8, Tumor) and FunCN(CD4, Tumor). These findings suggest that increased CAF activation and aggregation may lead to physical segregation between lymphocytes and tumor cells, thereby limiting immune cell infiltration, which was reflected in our simulations (Figure 4D, E, K). These findings are supported by samples with CAF activation marker having immunosuppressive microenvironment^34^. In addition to CAFs, the number of tumor-specific T cell clones (*n_T1_clones*) and the antigen binding rate (*k_P1_d1*) were positively associated with CD8+ T cell infiltration, indicating the importance of tumor-specific immunity.

**Fig. 7:**
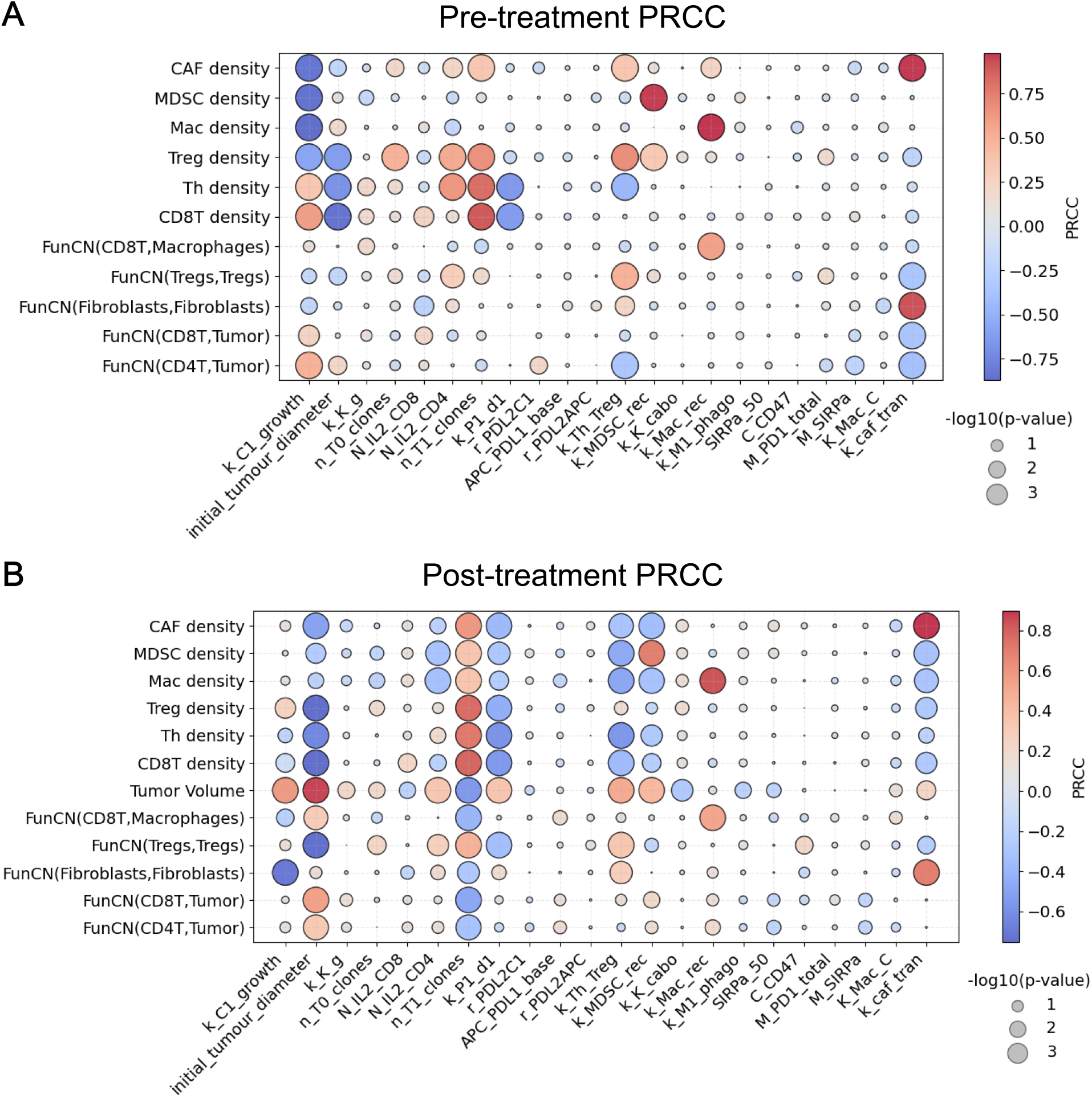
Partial Rank Correlation Coefficient (PRCC) sensitivity analysis for model parameters. **(A)** Pre-treatment and **(B)** post-treatment PRCC heatmaps showing the correlation between model parameters and selected spatial and non-spatial metrics. The color of each circle represents the value and direction of the PRCC (ρ), with red indicating positive and blue indicating negative correlation. The size of each circle reflects the statistical significance of the correlation, represented by the –log10 of the adjusted p-value (larger circles indicates more statistically significant associations).

In the post-treatment analysis, tumor growth rate (*k_C1_growth*), MDSC recruitment rate (*k_MDSC_rec*), and Treg transition rate (*k_Th_Treg*) were all positively correlated with tumor volume, reflecting their roles in promoting immunosuppression and therapeutic resistance.

Conversely, higher values of *n_T1_clones* were associated with smaller tumor volumes, emphasizing the critical role of tumor-specific T cell infiltration in mediating treatment response. Lastly, FunCN(Fibroblast, Fibroblast) remained positively correlated with both *k_caf_tran* and *k_Th_Treg*, suggesting that interactions between Tregs and CAFs contribute to fibroblast aggregation and spatial organization in the TME.

## Discussion

In this paper, we develop a computational pipeline to enable multi-scale spQSP modeling not only in simulating treatment dynamics but also in identifying spatial biomarkers for patient stratification and therapeutic decision-making^21,38^. A key highlight of this study is the use of pyABC^26^ to estimate the posterior distributions of cell motility parameters from a cohort of spatial molecular data representing static snapshots of the TME organization^26^. While demonstrated for an spQSP model, this approach can be generalized to the intractable likelihood functions commonly encountered in agent-based models (ABMs). Posterior estimation is performed through iterative simulations, where parameter sets are accepted based on simulated data that closely match observed data, measured by user-defined distance metrics. This approach allows us to calibrate complex multi-scale models based on available patient datasets, using spatial summary statistics to assimilate measurements such as spatial neighborhoods to infer model parameters.

Computationally, our modeling and calibration workflow has certain limitations. First, the design of an effective distance metric is critical, as it must balance the tradeoff between accurately capturing key features of the observed data and ensuring efficient convergence of the parameter estimation process. Second, the iterative nature of the framework can be computationally intensive, particularly when the distance metric is suboptimal, as this may lead to low acceptance rates of selected parameter sets and require large numbers of simulations. There are several alternative methods in the field, including Bayesflow^39^, Engine for Likelihood-Free Inference (ELFI)^40^, and PyTorch-based Simulation-Based Inference library (sbi)^41^. BayesFlow is an amortized, neural network-based framework for likelihood-free Bayesian inference that learns to map observed data to posterior distributions of model parameters using simulations ^39^.

Advantages of Bayesflow include scalability to high-dimensional data, compatibility with any model framework, and the ability to capture complex posteriors. Although the inference time of parameter estimation is fast due to its neural network-based approach, the upfront computational cost for training the neural network remains high. Engine for Likelihood-Free Inference (ELFI), a graph-based algorithm, allows data reuse parallelization, and visualization of inference workflows, which is suitable for computationally expensive model^40^. Finally, Simulation based Bayesian inference (SBI), a PyTorch-based toolbox, supports a variety of inference algorithms, neural network architectures, and sampling strategies, and includes built-in diagnostics and analysis tools for evaluating inference quality^41^.

In this case, we apply our computational framework to an spQSP model previously calibrated to HCC immunotherapy^6,7^. Our study confirms previous observations in a limited cohort of spatial transcriptomics data regarding the role of fibroblasts in blocking immune infiltration^24^ in a larger cohort of CODEX data. Based on these biological insights, this study also expanded our framework by incorporating a fibroblast module. Future work must extend our calibration pipeline to delineate when parameterization is sufficient, or new model components such as the addition of fibroblasts are needed to accurately reflect tumor biology. Still, through quantitative calibration of this extended spQSP model against pre-treatment spatial metrics derived from human biospecimens of HCC, the parametrized model captures key features of the tumor microenvironment. By simulating a virtual clinical trial, we stratified responders and non-responders and validated post-treatment predictions using FunCN scores from independent spatial proteomic and transcriptomic datasets. The model recapitulated clinically observed tumor architectures, quantified by the increased aggregation of fibroblasts in non-responders, and non-spatial features including higher Teff density and Teff/Treg ratio in responders. These results suggest our model accurately reflects spatial remodeling in the TME under combination therapy with nivolumab and cabozantinib. Additionally, our framework predicted response rates that closely matched clinical trials (CheckMate 040 and 459), demonstrating strong predictive fidelity. Although the response criteria in phase 3 trial (RECIST 1.1) differs from phase 1b trial^21^, the simulated response rates closely match the observed clinical response rates, demonstrating the robustness of our model. Importantly, the calibrated model enabled prospective biomarker discovery: spatial colocalization scores such as FunCN(CD8, Tumor) and FunCN(CD4, Tumor) emerged as positive prognostic indicators, while FunCN(Fibroblast, Fibroblast), FunCN(Treg, Treg), and FunCN(CD8, Macrophage) were associated with poor outcomes. Together, these findings highlight the utility of spQSP modeling not only in simulating treatment dynamics but also in identifying spatial biomarkers for patient stratification and therapeutic decision-making.

One limitation in this study is the fibroblast module, specifically that CAF aggregation difference is not significant at population level (Fig. 6H, I). Rather, the immunosuppressive effect of CAFs was observed at individual level (Fig. 6J-M, Supplemental Fig. 11). We attribute this discrepancy between simulation and clinical findings to potential sampling bias: non-responder specimens tend to be more tumor-dominant, whereas responder samples often contain more stromal regions, likely due to cancer cell elimination and concurrent ECM remodeling. This observation underscores the importance of distinguishing whether CAF accumulation is primarily driven by tumor-derived signaling or arises because of tissue repair mechanisms following tumor regression. Furthermore, although CAFs are known to be heterogeneous, encompassing subtypes such as myofibroblastic CAFs (myCAFs), inflammatory CAFs (iCAFs), and antigen-presenting CAFs (apCAFs) with distinct biological functions^38^, our omics data primarily revealed ECM remodeling that impedes immune cell–cancer cell interactions—a hallmark function of myCAFs. As such, our current model includes only the myCAF subtype. Additional CAF subtypes should be incorporated in the future as their functional roles become evident in our data. In addition, due to the widespread role of CAFs in immunosuppression future work must also evaluate how specific this model is to HCC applications or whether calibration against similar patient datasets in other tumor types would enable its broad applicability to immunotherapy response pan-cancer.

Despite the fact that we have successfully calibrated the spatial model using quantitative metrics, it is important to note that the calibration was performed at the population level using the full extensive patient cohort of CODEX data from Qiu et al^25^. There is no one-to-one correspondence between individual pre-treatment CODEX samples and the virtual patients generated in our simulations. This limitation also applies to our use of post-treatment IMC and Visium datasets, which were used for validation at cohort level rather than direct patient-level matching. The primary challenge preventing patient-specific simulations is due to the lack of paired longitudinal samples and challenges of obtaining temporal spatial molecular data due to cost and the invasive nature of the collection procedure. Specifically, this current study had the limitation that the pre-treatment and post-treatment datasets originate from different patient cohorts, as surgical resection is performed only once, limiting the availability of matched tissue samples across treatment timepoints. However, we acknowledge that individual-specific simulations, commonly referred to as digital twins, are essential for enabling personalized clinical decision-making.

Future work should complement these modeling strategies with digital twins fitted with longitudinal, patient-specific data, typically imaging data like MRI and CT scan to enhance data availability^42,43^. As data-driven artificial intelligence (AI) methods are gaining predictive power, generative cohorts of spatial molecular data may also overcome the limitations of sample profiling needed for model parameterization. Already, synthetic spatial images with varied cellular arrangements in the tumor microenvironment can simulate how spatial organization influences immune–tumor interactions, enabling hypothesis-driven exploration of factors affecting therapeutic efficacy^44–46^. Both model-driven and data-driven approaches have promises for allowing patient-specific simulations, comparing treatment outcomes across various therapeutic combinations and dosing regimens. In contrast to the mechanistic spQSP model, these data-driven AI models require minimal mechanistic understanding of underlying biology and may have comparable predictive power to the mechanistic model. However, they are more data-intensive and dependent on large, high-quality datasets and they lack the mechanistic interpretability of their mathematical modeling counterparts^44–46^. Therefore, future work must also cross-compare mechanistic and AI predictive models and develop hybrid approaches to best leverage the benefits of each strategy. In summary, the proposed data assimilation approach, which formally integrates patient-specific omics data into computational models, holds the potential to extend these models beyond virtual cohorts toward accurate outcome prediction at the individual patient level.

## Methods

### Spatial Quantitative System Pharmacology (spQSP) Model

The spQSP consists of a whole-body QSP model based on ordinary differential equations (ODEs) and an agent-based model (ABM) that captures the spatial structure of the tumor microenvironment at tissue level with a single-cell resolution. The integration method between the QSP and ABM components is detailed in the supplemental material. Mathematical formulations for previously developed modules are provided in the Supplemental Appendix, while the main text highlights new modules introduced in this study—specifically, modules for cabozantinib treatment, fibroblast interactions, and chemotaxis dynamics. Model parameters are available in the Supplemental Table. The full C++ source code for the model is publicly accessible, as outlined in the Data Availability Statement.

### Fibroblast Module

To model the biological function of fibroblasts in hepatocellular carcinoma (HCC) (Fig. 1A), we built upon prior fibroblast models developed for liver tissue^47^ and incorporated insights from our in-house Visium spatial transcriptomics data^24^. In the model, transforming growth factor-beta (TGFβ), secreted by immunosuppressive cells such as M2-like macrophages, cancer stem cells, and regulatory T cells (Tregs), induces the differentiation of hepatic stellate cells (HSCs) into cancer-associated fibroblasts (CAFs), which is modeled as:

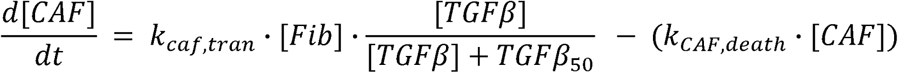

Here, _caf,tran_ is the transformation rate constant for fibroblasts converting into CAFs under the influence of TGFβ, which is calibrated based on pre-treatment fibroblast density from an _50_ is the half-maximal effective concentration of TGFβ. TGFβ in the tumor compartment.

### Extracellular Matrix (ECM) Modeling

In our previous spatial transcriptomic samples^24^, we observed that dense fibrotic regions act as physical barriers, obstructing direct contact between immune cells and cancer cells. To model this phenomenon, we formulated ECM secretion and degradation dynamics as follows:

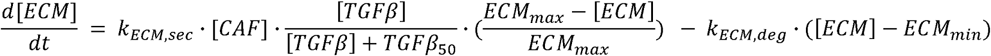

Here, and are ECM secretion and degradation rates by CAFs, respectively, _max_ and _min_ are maximum and minimum ECM density from Friedman et al.^47^. As previous studies have shown that ECM density negatively correlates with cell motility^49^, we modeled the ECM barrier effect on cell migration using the following equation:

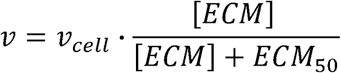

In this scenario, is the actual cell velocity governed by the local ECM density, and _cell_ is the reference cell velocity fitted from pre-treatment CODEX data^50^ (see Model Calibration and Validation section). _50_ is the half-maximal effective concentration of ECM that reduces cell motility by 50%.

### Cell Chemotaxis

While many studies have explored chemotaxis either in cancer contexts^51,52^ or within discrete cell-based frameworks^53^, few have addressed both areas simultaneously. To bridge this gap, we implement a run-and-reorient (i.e., run-and-tumble) scheme to simulate chemotactic cell behavior in response to cytokine gradients ^54,55^ . At each time step, a cell either continues moving in its current direction (“run”) or pauses to randomly select a new direction (“reorient,” analogous to tumbling). The probability of reorientation is modulated by the alignment between the cell’s current movement vector and the local chemokine gradient—cells are more likely to function of the cell’s velocity vector v·, the local cytokine concentration gradient 17,, and the

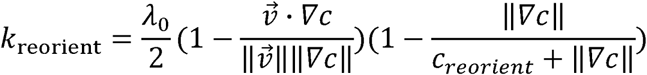

Here,10 denotes the maximum reorient rate, adapted from Friedman et al. ^47,52^. The term <u>^v^</u>represents the cosine of the angle between the cell’s velocity and the cytokine gradient, capturing the degree of directional alignment. The likelihood of a cell undergoing a directional change, representing a Poisson process. In our implementation, fibroblasts and regulatory T cells (Tregs) migrate along the TGFβ gradient; CD8 T cells and T helper (Th) cells follow the IFNγ gradient; and macrophages respond to CCL2 gradients.

### Cabozantinib Module

In this study, we retained the pharmacokinetic (PK) module of cabozantinib from our previously developed spQSP-HCC model and calibrated its pharmacodynamic (PD) module ^7^. Cabozantinib, a multi-tyrosine kinase inhibitor, exerts dual effects by inhibiting both cancer cell proliferation and tumor vasculature growth. The proliferation dynamics of cancer cells are modeled by the following differential equation:

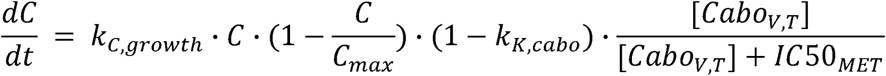

where C is the total number of cancer cells; Cmax is the carrying capacity of cancer cells in the parameters kC,growth and kK,cabo were calibrated using tumor growth data from a HepG2 section). The value for,C50MET is adopted from prior experimental measurements56. xenograft mouse model as reported in Xiang et al.28. (see Model Calibration using pyABC

### Spatial molecular datasets for analysis

To ensure that the spatial organization generated by the ABM accurately recapitulates the tumor microenvironments observed in clinical samples, we calibrated the simulation framework using spatial proteomics data (CODEX) from 401 treatment-naive HCC patients, as reported in Qiu et al.^25^. Cell segmentation and phenotyping were conducted in the original study, and we utilized the phenotyped results to compute FunCN scores. Post-treatment data are acquired from a previously conducted phase 1b clinical trial of cabozantinib plus nivolumab (CABO/NIVO), prospectively registered at ClinicalTrials.gov (NCT03299946). A dataset comprising 36 tissue microarrays (TMAs) from 12 HCC patients profiled by imaging mass cytometry (IMC) with 27 function markers from that clinical trial, generated in Ho et al.^21^ and annotated as described in Mi et al.^33^; cell segmentation and phenotyping were also included in the original study, similar to pre-treatment CODEX data. Additionally, we incorporated Visium spatial transcriptomics data from the same clinical trial^24^ to further support our findings. However, FunCN scoring was not applied to the Visium dataset due to its lack of single-cell resolution.

### Functional Cell Neighborhood (FunCN) Score

In this study, we used FunCN scores to quantitatively analyze both CODEX and IMC datasets. To quantify spatial influence of specific cell types (Source cell) around a given target cell, we apply Gaussian kernel density estimation^27^. Each neighboring cell contributes to the spatial weight based on its Euclidean distance to the target cell, modulated by a Gaussian kernel function:

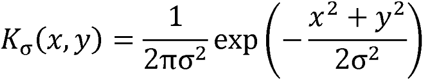

Here, equals 10 microns for both simulation and spatial-omics data, determining the spatial scale over which influence is considered. Points closer to the target cell receive higher weights, while more distant cells contribute less. For a target location, the total spatial weight for a particular reference cell type is computed by summing the kernel-weighted contributions from all cells of that type:

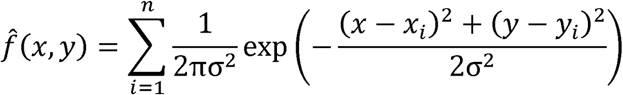

This gives a cell-cell-specific spatial neighborhood score reflecting the density-weighted proximity of the reference cells around each target cell, allowing quantitative comparisons between spatial proteomic data and model outputs. In results section, we use FunCN(A, B) to represent the average spatial influence of cell type B (Source cell) around cell type A (Target Cell).

### Model Calibration using pyABC

We employed pyABC framework^26^ to implement the Approximate Bayesian Computation - Sequential Monte Carlo (ABC-SMC) model calibration approach for model parameter inference which iterates model calibration over multiple populations^57^. For each population, particles representing parameter combination are first sampled from a prior distribution and passed to either QSP or spQSP model for simulation. Simulation outputs are compared with accepted to enter the next population in which the posterior distribution rr e y from the current population is used as an updated prior. Here, we used uniform distribution as the prior in the first population, with the calibration running for five populations. *minimum_epsilon*, the stopping criterion for adaptive rejection sampling, set to 0.01. All other parameters are set as default in this study. For non-spatial parameter calibration in the QSP, mouse tumor volume measured at different time points was used as the observational data^28^. For ABM parameter calibration, selected FunCN scores served as observational data, using the Euclidian distance between simulation FunCN score and observed FunCN score as distance metrics. Code availability for model calibration using pyABC is in the Data Availability Statement.

### Statistical Analysis

We used log-ranked test for comparing overall survival between high-FunCN vs. low-FunCN group. Computed p-values are adjusted by false discovery rate (FDR). For comparing FunCN score and cell densities between responder versus non-responder, we used Wilcoxon ranksum test. Finally, for partial ranked correlation coefficient (PRCC), we used t-test for correlation coefficients ρ. These tests are either performed in MATLAB or R, depending on the simulation software.

### Sensitivity Analysis

To assess the influence of model parameters on both spatial and non-spatial response metrics in virtual patients, including FunCN scores and cell densities, we performed sensitivity analysis on 18 parameters for sampling virtual patients using Partial Rank Correlation Coefficient (PRCC) analysis. Parameter values are sampled with the LHS method, and distributions are shown in supplemental tables.

## Funding

This work was funded by NIH U24CA284156 to EJF and AD, U01CA253403 to EJF, and U01CA212007 to EJF and ASP. Spatial transcriptomic analysis was supported by DoD- MEDCOM 144517–CA220654P to LTK. This work was made possible in part through the support of the Maryland Cancer Moonshot Research Grant to the Johns Hopkins Medical Institutions (FY24) to EJF and AD, and the Maryland Cigarette Restitution fund (FY25) to AD. Figure 1 was created with BioRender.com.

## Data Availability Statement

The authors confirm that the data supporting the findings of this study are available within the article and the Supplement. C++ code for model generation and virtual clinical trials will be available upon acceptance. The pyABC model calibration code is available at will also be available upon acceptance

## Conflicts of Interest

M. Yarchoan reports grants and personal fees from Genentech, Exelixis, and AstraZeneca, grants from Bristol Myers Squibb and Incyte, personal fees from Replimune, Hepion, and Lantheus, and other support from Adventris outside the submitted work. E.M. Jaffee reports other support from Abmeta and Adventris, personal fees from Dragonfly, Neuvogen, STIMIT Tx, Mestag, Candel therapeutics, HDTbio, and grants from Lustgarten, Genentech, BMS, NeoTx, and Break Through Cancer. Dr. Elizabeth Jaffee is a founder of and holds equity in Adventris Pharmaceuticals. She also serves as a consultant to the entity. W.J. Ho reports patent royalties from Rodeo/Amgen; completed grants from Sanofi and NeoTX; and speaking/travel honoraria from Exelixis and Standard BioTools. E.J. Fertig reports grants from NIH/NCI and Johns Hopkins University during the conduct of the study; personal fees from Mestag Therapeutics, Resistance Bio, and Merck; and grants from AbbVie, Inc., Roche/Genentech, Lustgarten Foundation, Break Through Cancer, Emerson Collective, and NIH outside the submitted work. A.S. Popel receives research funding from Merck. No disclosures were reported by the other authors. The terms of these arrangements are being managed by the Johns Hopkins University in accordance with its conflict-of-interest policies. The remaining authors declare that the research was conducted in the absence of any commercial or financial relationships that could be construed as a potential conflict of interest.

## References

1. Anbari S, Wang H, Arulraj T, et al. Identifying biomarkers for treatment of uveal melanoma by T cell engager using a QSP model. NPJ Syst Biol Appl. 2024;10(1):108. doi:10.1038/s41540-024-00434-5

2. Wang H, Arulraj T, Ippolito A, Popel AS. From virtual patients to digital twins in immuno-oncology: lessons learned from mechanistic quantitative systems pharmacology modeling. NPJ Digit Med. 2024;7(1):1–6. doi:10.1038/s41746-024-01188-4

3. Arulraj T, Wang H, Deshpande A, et al. Virtual patient analysis identifies strategies to improve the performance of predictive biomarkers for PD-1 blockade. Proc Natl Acad Sci. 2024;121(45):e2410911121. doi:10.1073/pnas.2410911121

4. Ippolito A, Wang H, Zhang Y, Vakil V, Popel AS. Virtual clinical trials via a QSP immuno-oncology model to simulate the response to a conditionally activated PD-L1 targeting antibody in NSCLC. J Pharmacokinet Pharmacodyn. Published online 2024:747–757. doi:10.1007/s10928-024-09928-5

5. Wang H, Zhao C, Santa-Maria CA, Emens LA, Popel AS. Dynamics of tumor-associated macrophages in a quantitative systems pharmacology model of immunotherapy in triple-negative breast cancer. iScience. 2022;25(8):104702. doi:10.1016/j.isci.2022.104702

6. Sové RJ, Verma BK, Wang H, Ho WJ, Yarchoan M, Popel AS. Virtual clinical trials of anti-PD-1 and anti-CTLA-4 immunotherapy in advanced hepatocellular carcinoma using a quantitative systems pharmacology model. J Immunother Cancer. 2022;10(11):e005414. doi:10.1136/jitc-2022-005414

7. Zhang S, Deshpande A, Verma BK, et al. Integration of Clinical Trial Spatial Multi-omics Analysis and Virtual Clinical Trials Enables Immunotherapy Response Prediction and Biomarker Discovery. Cancer Res. Published online 2024:2734–2748. doi:10.1158/0008-5472.can-24-0943

8. Gong C, Milberg O, Wang B, et al. A computational multiscale agent-based model for simulating spatio-temporal tumour immune response to PD1 and PDL1 inhibition. J R Soc Interface. 2017;14(134). doi:10.1098/rsif.2017.0320

9. Ruiz-Martinez A, Gong C, Wang H, et al. Simulations of tumor growth and response to immunotherapy by coupling a spatial agent-based model with a whole-patient quantitative systems pharmacology model. PLoS Comput Biol. 2022;18(7):1–32. doi:10.1371/journal.pcbi.1010254

10. Zhang S, Gong C, Ruiz-Martinez A, et al. Integrating single cell sequencing with a spatial quantitative systems pharmacology model spQSP for personalized prediction of triple-negative breast cancer immunotherapy response. ImmunoInformatics. 2021;1–2(July):100002. doi:10.1016/j.immuno.2021.100002

11. Mi H, Sivagnanam S, Ho WJ, et al. Computational methods and biomarker discovery strategies for spatial proteomics: a review in immuno-oncology. Brief Bioinform. 2024;25(5). doi:10.1093/bib/bbae421

12. Johnson JAI, Bergman DR, Rocha HL, et al. Human interpretable grammar encodes multicellular systems biology models to democratize virtual cell laboratories. Cell. 2025;188(17):4711–4733.e37. doi:10.1016/j.cell.2025.06.048

13. Cess CG, Finley SD. Calibrating agent-based models to tumor images using representation learning. PLoS Comput Biol. 2023;19(4):e1011070. doi:10.1371/journal.pcbi.1011070

14. Kostelich EJ, Kuang Y, Mcdaniel JM, Moore NZ, Martirosyan NL, Preul MC. Accurate state estimation from uncertain data and models : an application of data assimilation to mathematical models of human brain tumors. Published online 2011:1–20. doi:10.1186/1745-6150-6-64.

15. Siegel RL, Kratzer TB, Giaquinto AN, Sung H, Jemal A. Cancer statistics, 2025. CA Cancer J Clin. 2025;(October 2024):10-45. doi:10.3322/caac.21871

16. Singal AG, Kanwal F, Llovet JM. Global trends in hepatocellular carcinoma epidemiology: implications for screening, prevention and therapy. Nat Rev Clin Oncol. 2023;20(12):864–884. doi:10.1038/s41571-023-00825-3

17. Finn RS, Kudo M, Merle P, et al. LBA34 Primary results from the phase III LEAP-002 study: Lenvatinib plus pembrolizumab versus lenvatinib as first-line (1L) therapy for advanced hepatocellular carcinoma (aHCC). Ann Oncol. 2022;33:S1401. doi:10.1016/j.annonc.2022.08.031

18. Qin S, Bi F, Gu S, et al. Donafenib Versus Sorafenib in First-Line Treatment of Unresectable or Metastatic Hepatocellular Carcinoma: A Randomized, Open-Label, Parallel-Controlled Phase II-III Trial. J Clin Oncol. 2021;39(27):3002–3011. doi:10.1200/JCO.21.00163

19. Yau T, Park JW, Finn RS, et al. CheckMate 459: A randomized, multi-center phase III study of nivolumab (NIVO) vs sorafenib (SOR) as first-line (1L) treatment in patients (pts) with advanced hepatocellular carcinoma (aHCC). Ann Oncol. 2019;30(October):v874–v875. doi:10.1093/annonc/mdz394.029

20. Yau T, Zagonel V, Santoro A, et al. Nivolumab Plus Cabozantinib with or Without Ipilimumab for Advanced Hepatocellular Carcinoma: Results from Cohort 6 of the CheckMate 040 Trial. J Clin Oncol. 2023;41(9):1747–1757. doi:10.1200/JCO.22.00972

21. Ho WJ, Zhu Q, Durham J, et al. Neoadjuvant cabozantinib and nivolumab convert locally advanced hepatocellular carcinoma into resectable disease with enhanced antitumor immunity. Nat Cancer. 2021;2(9):891–903. doi:10.1038/s43018-021-00234-4

22. Yarchoan M, Gane EJ, Marron TU, et al. Personalized neoantigen vaccine and pembrolizumab in advanced hepatocellular carcinoma: a phase 1/2 trial. Nat Med. 2024;30(4):1044–1053. doi:10.1038/s41591-024-02894-y

23. Greten TF, Villanueva A, Korangy F, et al. Biomarkers for immunotherapy of hepatocellular carcinoma. Nat Rev Clin Oncol. 2023;20(11):780–798. doi:10.1038/s41571-023-00816-4

24. Zhang S, Yuan L, Danilova L, et al. Spatial transcriptomics analysis of neoadjuvant cabozantinib and nivolumab in advanced hepatocellular carcinoma identifies independent mechanisms of resistance and recurrence. Genome Med. 2023;15(1):72. doi:10.1186/s13073-023-01218-y

25. Qiu X, Zhou T, Li S, et al. Spatial single-cell protein landscape reveals vimentinhigh macrophages as immune-suppressive in the microenvironment of hepatocellular carcinoma. Nat Cancer. 2024;5(October). doi:10.1038/s43018-024-00824-y

26. Schälte Y, Klinger E, Alamoudi E, Hasenauer J. pyABC: Efficient and robust easy-to-use approximate Bayesian computation. J Open Source Softw. 2022;7(74):4304. doi:10.21105/joss.04304

27. Cho Y, Lee JW, Shin SM, et al. Modeling cellular influence delineates functionally relevant cellular neighborhoods in primary and metastatic pancreatic ductal adenocarcinoma. *bioRxiv*. Published online January 1, 2025:2025.06.12.659314. doi:10.1101/2025.06.12.659314

28. Xiang Q, Chen W, Ren M, et al. Cabozantinib suppresses tumor growth and metastasis in hepatocellular carcinoma by a dual blockade of VEGFR2 and MET. Clin Cancer Res. 2014;20(11):2959–2970. doi:10.1158/1078-0432.CCR-13-2620

29. Reed DR, Bachmanov AA, Tordoff MG. Forty mouse strain survey of body composition. Physiol Behav. 2007;91(5):593–600. 10.1016/j.physbeh.2007.03.026

30. Abou-Alfa GK, Meyer T, Cheng A-L, et al. Cabozantinib in Patients with Advanced and Progressing Hepatocellular Carcinoma. N Engl J Med. 2018;379(1):54–63. doi:10.1056/nejmoa1717002

31. Sangro B, Sarobe P, Hervás-Stubbs S, Melero I. Advances in immunotherapy for hepatocellular carcinoma. Nat Rev Gastroenterol Hepatol. 2021;18(8):525–543. doi:10.1038/s41575-021-00438-0

32. Wang H, Liang Y, Liu Z, et al. POSTN+ cancer-associated fibroblasts determine the efficacy of immunotherapy in hepatocellular carcinoma. J Immunother Cancer. 2024;12(7). doi:10.1136/jitc-2023-008721

33. Mi H, Ho WJ, Yarchoan M, Popel AS. Multi-Scale Spatial Analysis of the Tumor Microenvironment Reveals Features of Cabozantinib and Nivolumab Efficacy in Hepatocellular Carcinoma. Front Immunol. 2022;13(May):1–16. doi:10.3389/fimmu.2022.892250

34. Xue R, Zhang Q, Cao Q, et al. Liver tumour immune microenvironment subtypes and neutrophil heterogeneity. Nature. 2022;612(7938):141-147. doi:10.1038/s41586-022-05400-x

35. Ahn B, Ahn H-S, Shin J, et al. Characterization of lymphocyte-rich hepatocellular carcinoma and the prognostic role of tertiary lymphoid structures. Liver Int. 2024;44(5):1202–1218. 10.1111/liv.15865

36. Rodriguez AB, Peske JD, Woods AN, et al. Immune mechanisms orchestrate tertiary lymphoid structures in tumors via cancer-associated fibroblasts. Cell Rep. 2021;36(3):109422. doi:10.1016/j.celrep.2021.109422

37. Liu Y, Xun Z, Ma K, et al. Identification of a tumour immune barrier in the HCC microenvironment that determines the efficacy of immunotherapy. J Hepatol. 2023;78(4):770–782. doi:10.1016/j.jhep.2023.01.011

38. Yang D, Liu J, Qian H, Zhuang Q. Cancer-associated fibroblasts: from basic science to anticancer therapy. Exp Mol Med. 2023;55(7):1322–1332. doi:10.1038/s12276-023-01013-0

39. Radev ST, Mertens UK, Voss A, Ardizzone L, Kothe U. BayesFlow: Learning Complex Stochastic Models With Invertible Neural Networks. IEEE Trans Neural Networks Learn Syst. 2022;33(4):1452–1466. doi:10.1109/TNNLS.2020.3042395

40. Lintusaari J, Vuollekoski H, Kangasrääsiö A, et al. ELFI: Engine for likelihood-free inference. J Mach Learn Res. 2018;19:1–7.

41. Boelts J, Deistler M, Gloeckler M, et al. sbi reloaded : a toolkit for simulation-based inference workflows. J Open Source Softw. 2025;10:1–9. doi:10.21105/joss.07754

42. Hormuth DA, Eldridge SL, Weis JA, Miga MI, Yankeelov TE. Mechanically Coupled Reaction-Diffusion Model to Predict Glioma Growth: Methodological Details BT - Cancer Systems Biology: Methods and Protocols. In: von Stechow L, ed. Springer New York; 2018:225–241. doi:10.1007/978-1-4939-7493-1_11

43. Jarrett AM, Kazerouni AS, Wu C, et al. Quantitative magnetic resonance imaging and tumor forecasting of breast cancer patients in the community setting. Nat Protoc. 2021;16(11):5309–5338. doi:10.1038/s41596-021-00617-y

44. Quiros AC, Murray-Smith R, Yuan K. PathologyGAN: Learning deep representations of cancer tissue. Proc Mach Learn Res. 2020;121(Midl):669–695. doi:10.59275/j.melba.2021-gfgg

45. Moghadam PA, Van Dalen S, Martin KC, et al. A Morphology Focused Diffusion Probabilistic Model for Synthesis of Histopathology Images. Proc - 2023 *IEEE Winter Conf Appl Comput Vision, WACV* 2023. Published online 2023:1999-2008. doi:10.1109/WACV56688.2023.00204

46. Kazerouni A, Aghdam EK, Heidari M, et al. Diffusion models in medical imaging: A comprehensive survey. Med Image Anal. 2023;88(November 2022):102846. doi:10.1016/j.media.2023.102846

47. Friedman A, Hao W. Mathematical modeling of liver fibrosis. Math Biosci Eng. 2017;14(1):143–164. doi:10.3934/mbe.2017010

48. Ju MJ, Qiu SJ, Fan J, et al. Peritumoral activated hepatic stellate cells predict poor clinical outcome in hepatocellular carcinoma after curative resection. Am J Clin Pathol. 2009;131(4):498–510. doi:10.1309/AJCP86PPBNGOHNNL

49. Salmon H, Franciszkiewicz K, Damotte D, et al. Matrix architecture defines the preferential localization and migration of T cells into the stroma of human lung tumors. J Clin Invest. 2012;122(3):899–910. doi:10.1172/JCI45817

50. Qiu X, Zhou T, Li S, et al. Spatial single-cell protein landscape reveals vimentinhigh macrophages as immune-suppressive in the microenvironment of hepatocellular carcinoma. Nat Cancer. 2024;5(10):1557–1578. doi:10.1038/s43018-024-00824-y

51. Mpekris F, Voutouri C, Baish JW, et al. Combining microenvironment normalization strategies to improve cancer immunotherapy. Proc Natl Acad Sci U S A. 2020;117(7):3728–3737. doi:10.1073/pnas.1919764117

52. Kim Y, Friedman A. Interaction of Tumor with Its Micro-environment: A Mathematical Model. Bull Math Biol. 2010;72(5):1029–1068. doi:10.1007/s11538-009-9481-z

53. Milde F, Bergdorf M, Koumoutsakos P. A hybrid model for three-dimensional simulations of sprouting angiogenesis. Biophys J. 2008;95(7):3146–3160. doi:10.1529/biophysj.107.124511

54. Nakamura K, Kobayashi TJ. Optimal sensing and control of run-and-tumble chemotaxis. Phys Rev Res. 2022;4(1):4–6. doi:10.1103/PhysRevResearch.4.013120

55. Santra I, Basu U, Sabhapandit S. Run-and-tumble particles in two dimensions: Marginal position distributions. Phys Rev E. 2020;101(6):1–19. doi:10.1103/PhysRevE.101.062120

56. Osanto S, van der Hulle T. Cabozantinib in the treatment of advanced renal cell carcinoma in adults following prior vascular endothelial growth factor targeted therapy: clinical trial evidence and experience. Ther Adv Urol. 2018;10(3):109–123. doi:10.1177/1756287217748867

57. Toni T, Stumpf MPH. Simulation-based model selection for dynamical systems in systems and population biology. Bioinformatics. 2010;26(1):104–110. doi:10.1093/bioinformatics/btp619

