## Supplemental Figure for "Quantitative Calibration of a Spatial QSP Model Identifies Fibroblast Impact on HCC Immunotherapy"

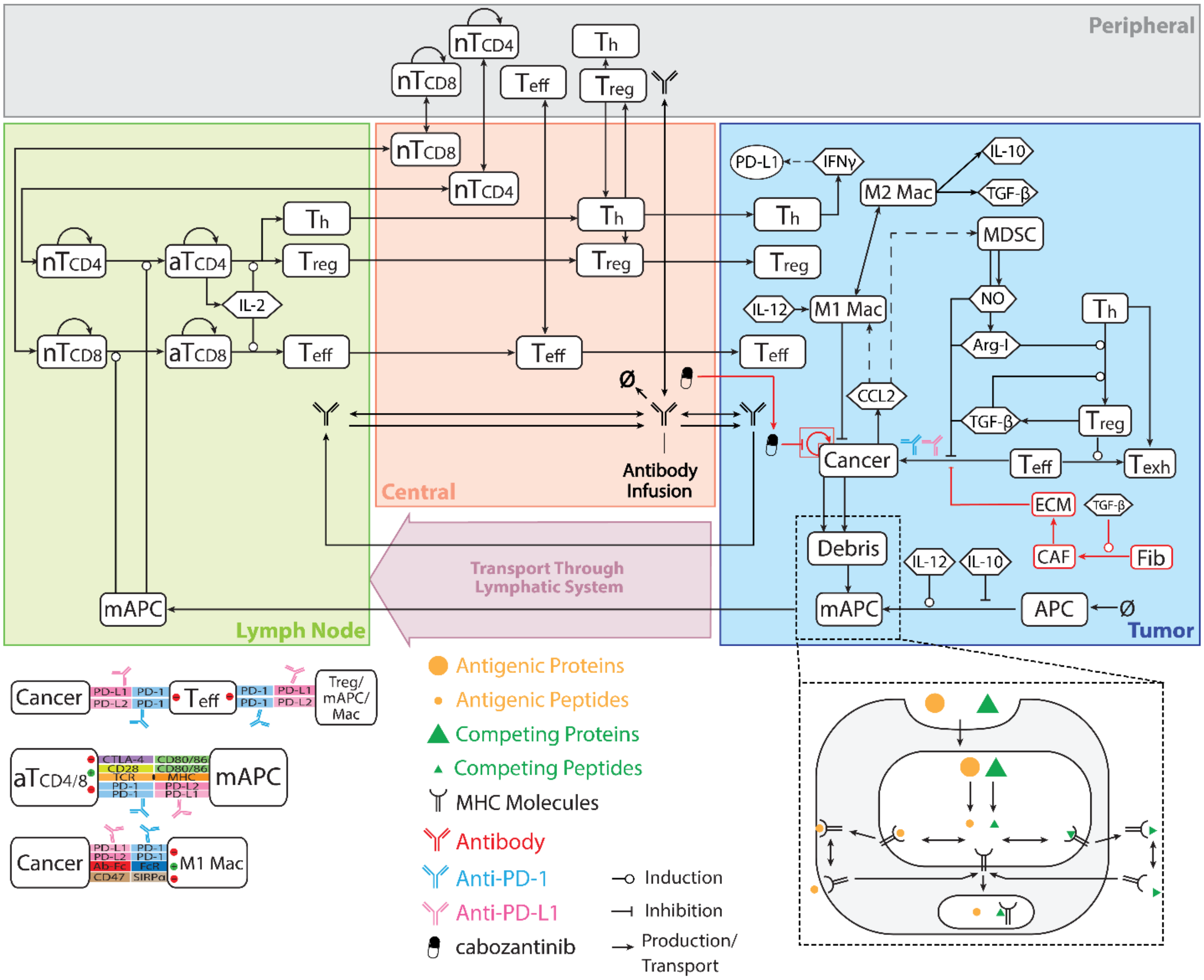

**Supplemental Fig. 1** Whole-body QSP model simulating immune dynamics, antigen presentation, and pharmacokinetics/pharmacodynamics (PK/PD) of selected therapies. Newly added modules are highlighted in red.

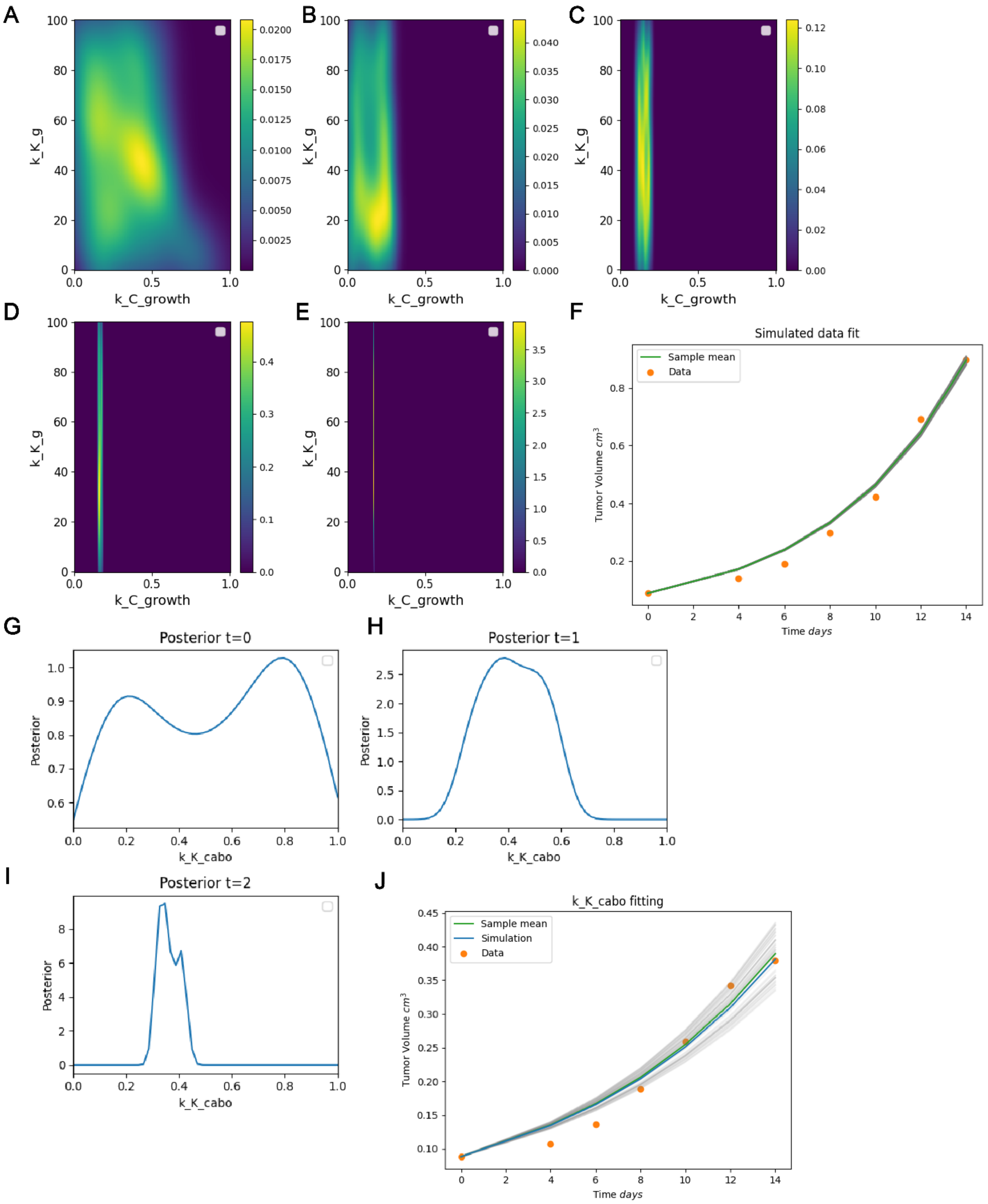


Supplemental Fig. 2: Cabozantinib module recalibration result using mouse model from Xiang et al. **(A-E)** Two-dimensional heatmap showing posterior distribution of tumor vasculature growth rate (*k_K_g*) and tumor growth rate (*k_C_growth*) over 5 iterations using pyABC. **(F)** Simulated tumor growth trajectories after calibration. Ten parameter sets were sampled from the final posterior distribution; the green line represents the simulation mean, and the orange dots represent the experimental data. (G-I) Density plot showing posterior distribution of vasculature and tumor growth inhibition effect of cabozantinib (*k_K_cabo*) over three pyABC iterations. **(J)** Tumor growth fitting using the calibrated *k_K_cabo*. Gray lines indicate individual simulation trajectories; the green line shows the mean; orange dots indicate experimental data.

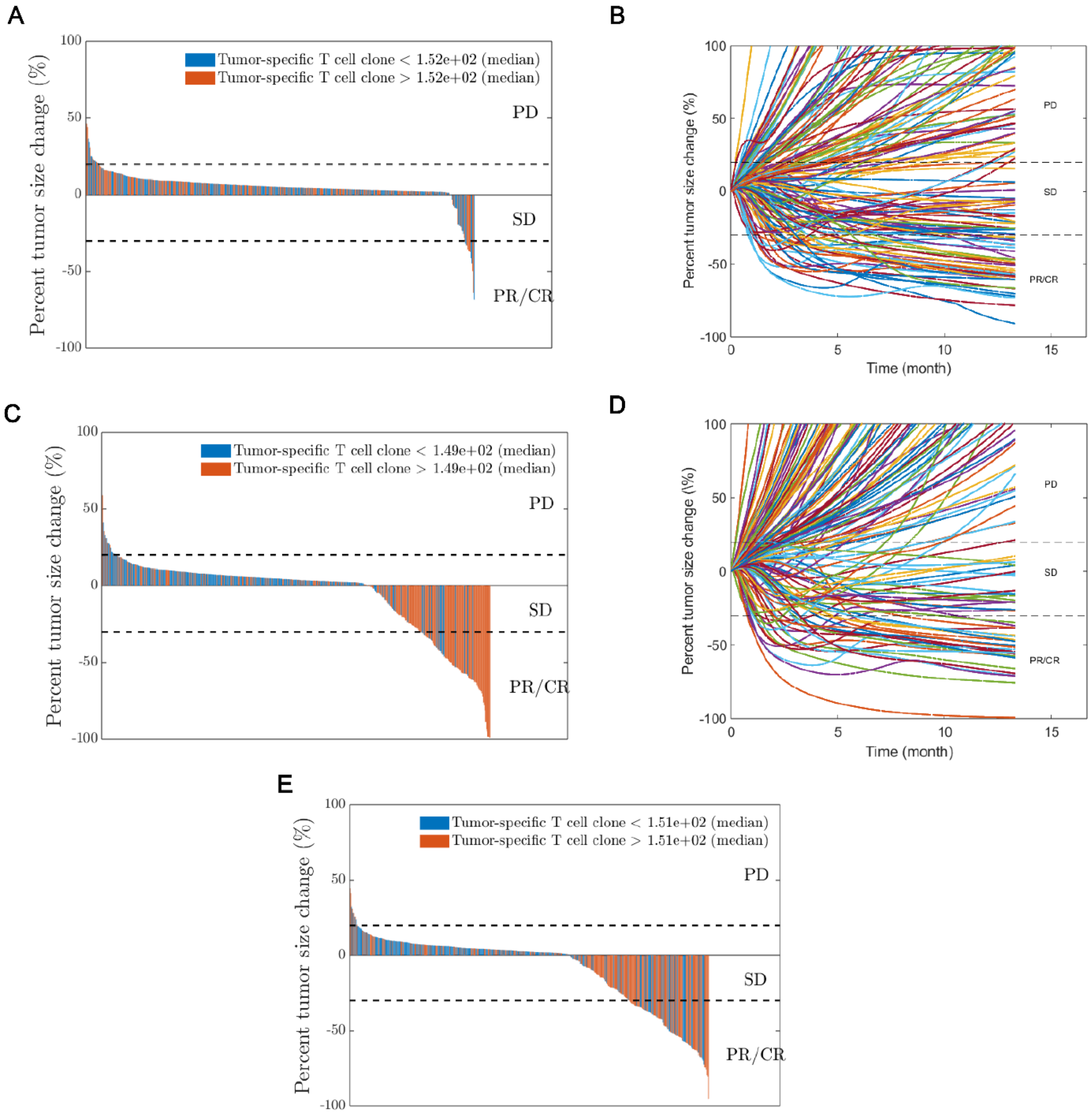


Supplemental Fig 3. QSP model calibration using single-arm clinical trial data. **(A)** Waterfall plot of simulated tumor size changes in a virtual clinical trial of cabozantinib monotherapy, calibrated to the Abou-Alfa et al. trial. **(B)** Spider plot showing individual tumor trajectories over time for virtual patients treated with cabozantinib monotherapy for 150 randomly selected virtual patients out of 500. Tumor responses are categorized as progressive disease (PD), stable disease (SD), or partial/complete response (PR/CR) according to RECIST-like criteria. **(C)** Waterfall plot of simulated tumor size changes in a virtual clinical trial of nivolumab monotherapy, calibrated to the CheckMate 459 trial. Patients are stratified by the median number of tumor-specific T cell clones. **(D)** Spider plot showing individual tumor trajectories over time for virtual patients treated with nivolumab monotherapy for 150 randomly selected virtual patients out of 500. **(E)** Waterfall plot of simulated tumor size changes in a virtual clinical trial of nivolumab and cabozantinib for model validation, color-coded by tumor-specific T cell clone counts: above (orange) or below (blue) the median of tumor specific clones (n_T1_clone).

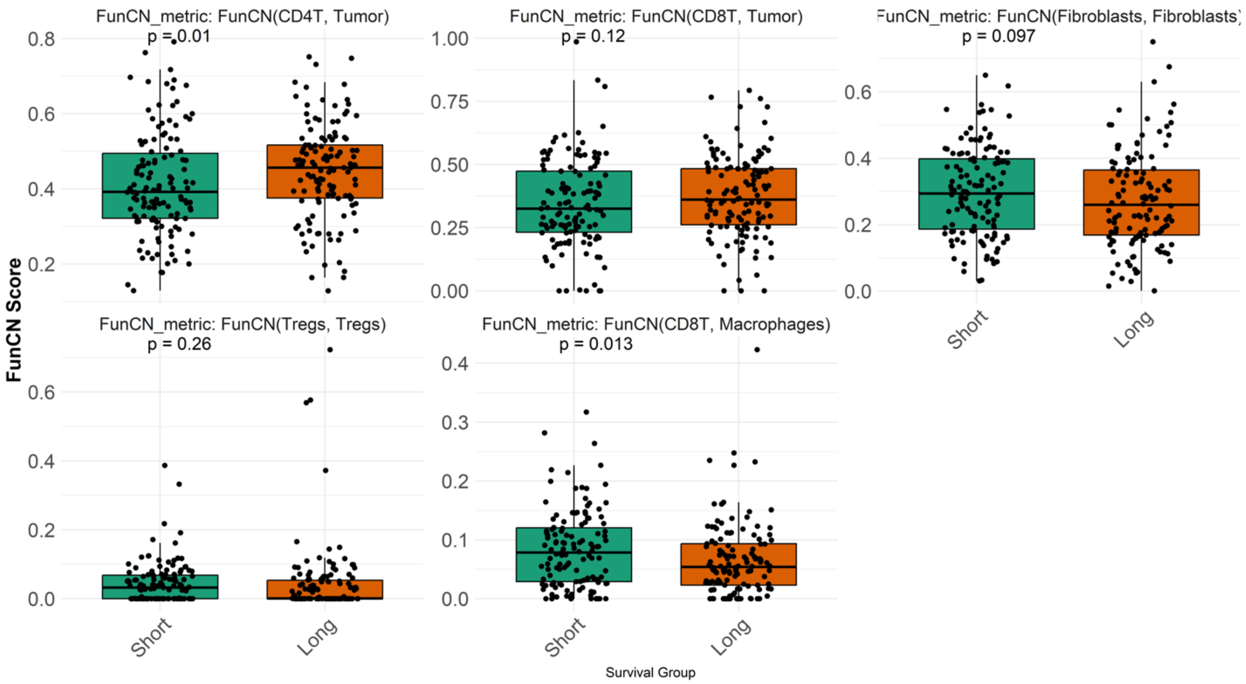


Supplemental Fig. 4: Boxplots of five post-treatment FunCN scores stratified by overall survival spilt by median OS days (Long term vs. Short term). P-values indicate statistical significance for each metric.

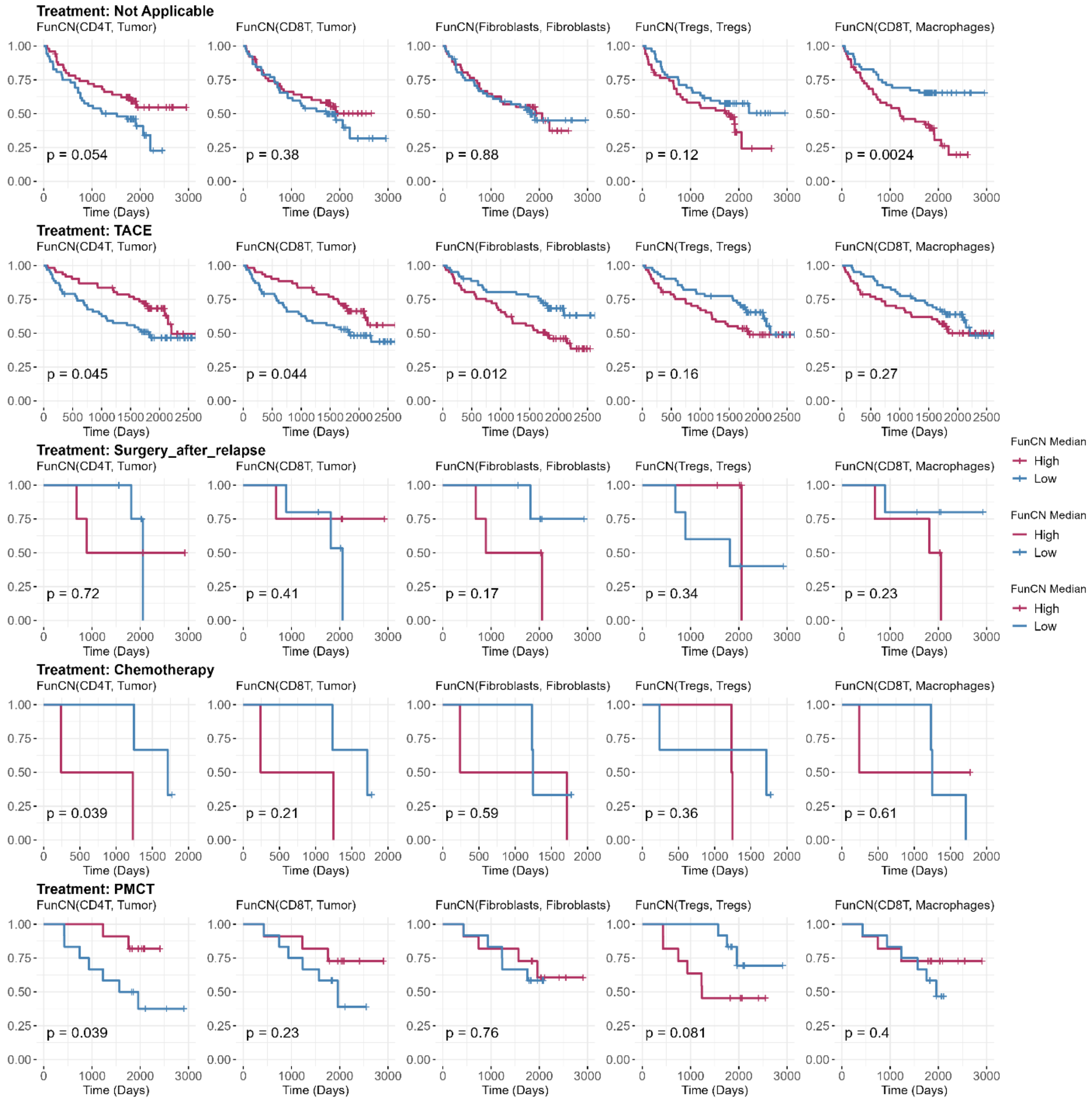


Supplemental Fig. 5. Kaplan–Meier survival curves stratified by five selected FunCN metrics across different post-surgical treatment groups. Each panel compares patient cohorts with high versus low FunCN scores (spilt at the median). Statistical significance was assessed using the log-rank test, with p-values adjusted by the False Discovery Rate (FDR).

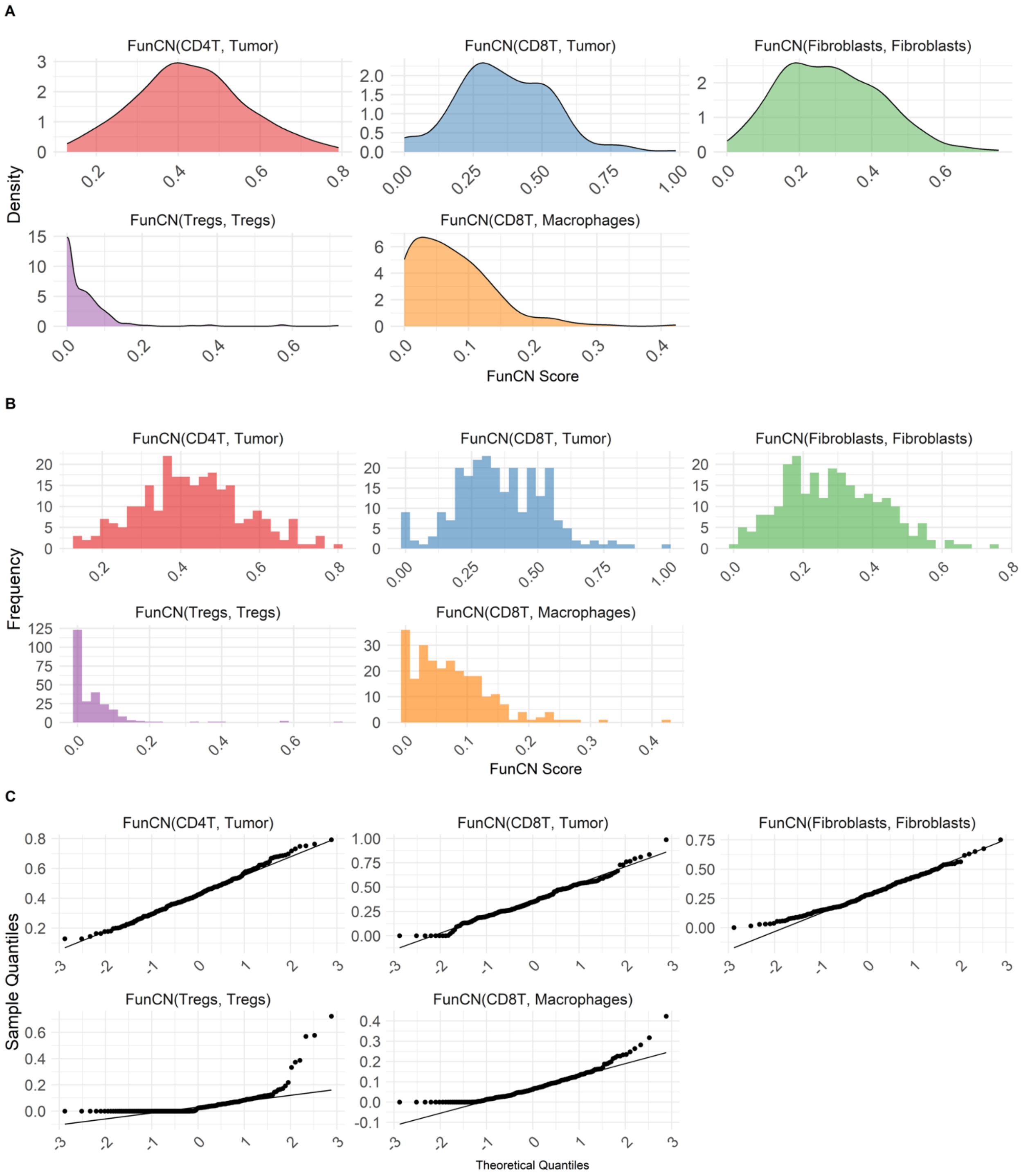


Supplemental Fig. 6: Distribution analysis of five pre-treatment FunCN scores derived from the CODEX dataset (Qiu et al). **(A)** Density plots showing the distribution of FunCN scores for CD4$^{+}$ T cells–tumor, CD8$^{+}$ T cells–tumor, fibroblast–fibroblast, Treg–Treg, and CD8$^{+}$ T cells–macrophage interactions. **(B)** Corresponding frequency histograms for each FunCN score. **(C)** Q–Q plots assessing the normality of each FunCN score distribution by comparing sample quantiles to theoretical normal quantiles.

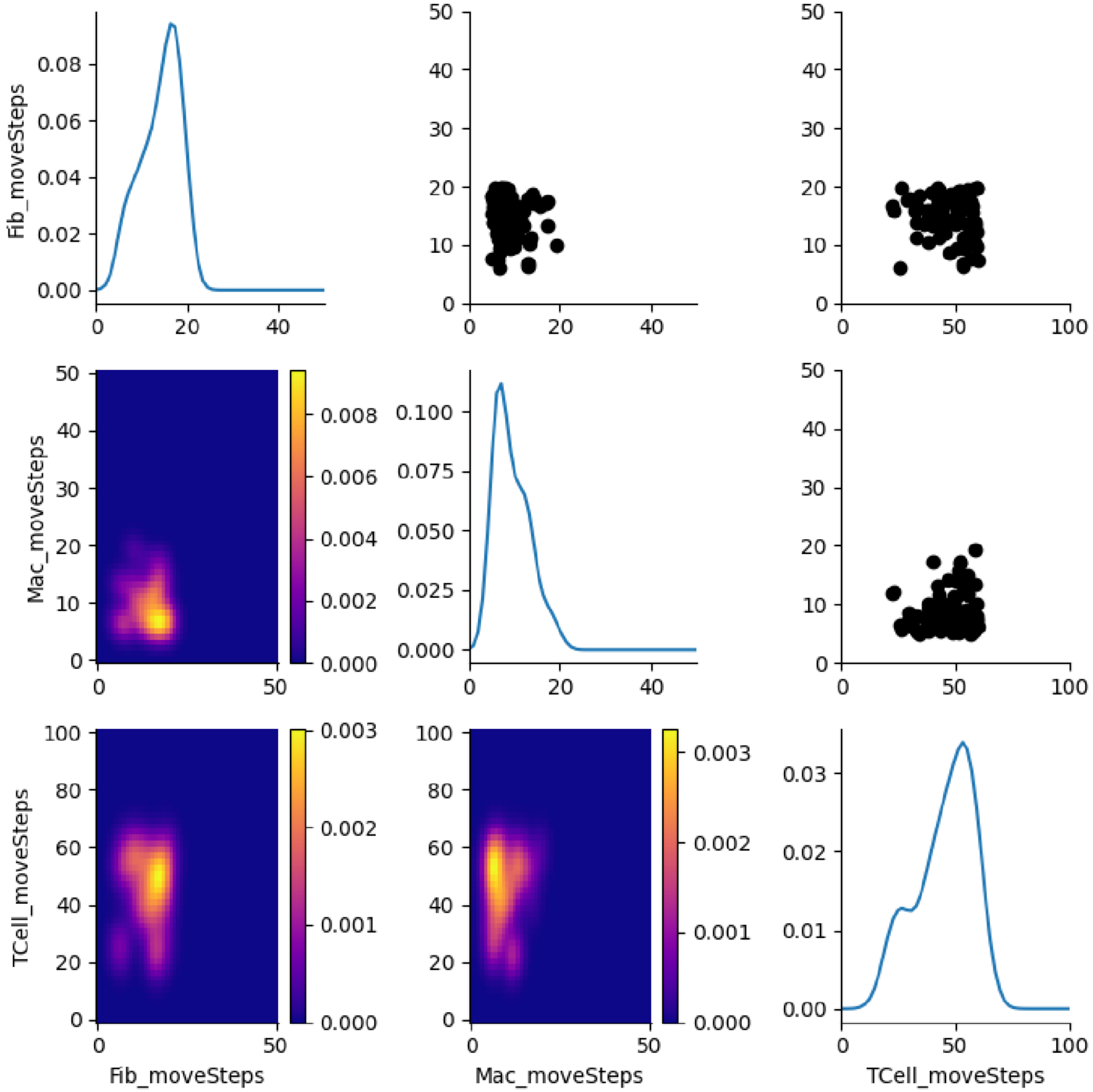


Supplemental Fig. 7: Calibration of immune and stromal cell motility parameters based on pre-treatment FunCN scores**.** Diagonal plots show the posterior distributions of cell motilities for fibroblasts, macrophages, and T cells after pyABC calibration. Plots in the upper right triangle represent accepted parameter instances, where each dot corresponds to a sampled parameter combination. Plots in the lower left triangle display two-dimensional heatmaps of the joint posterior distributions.

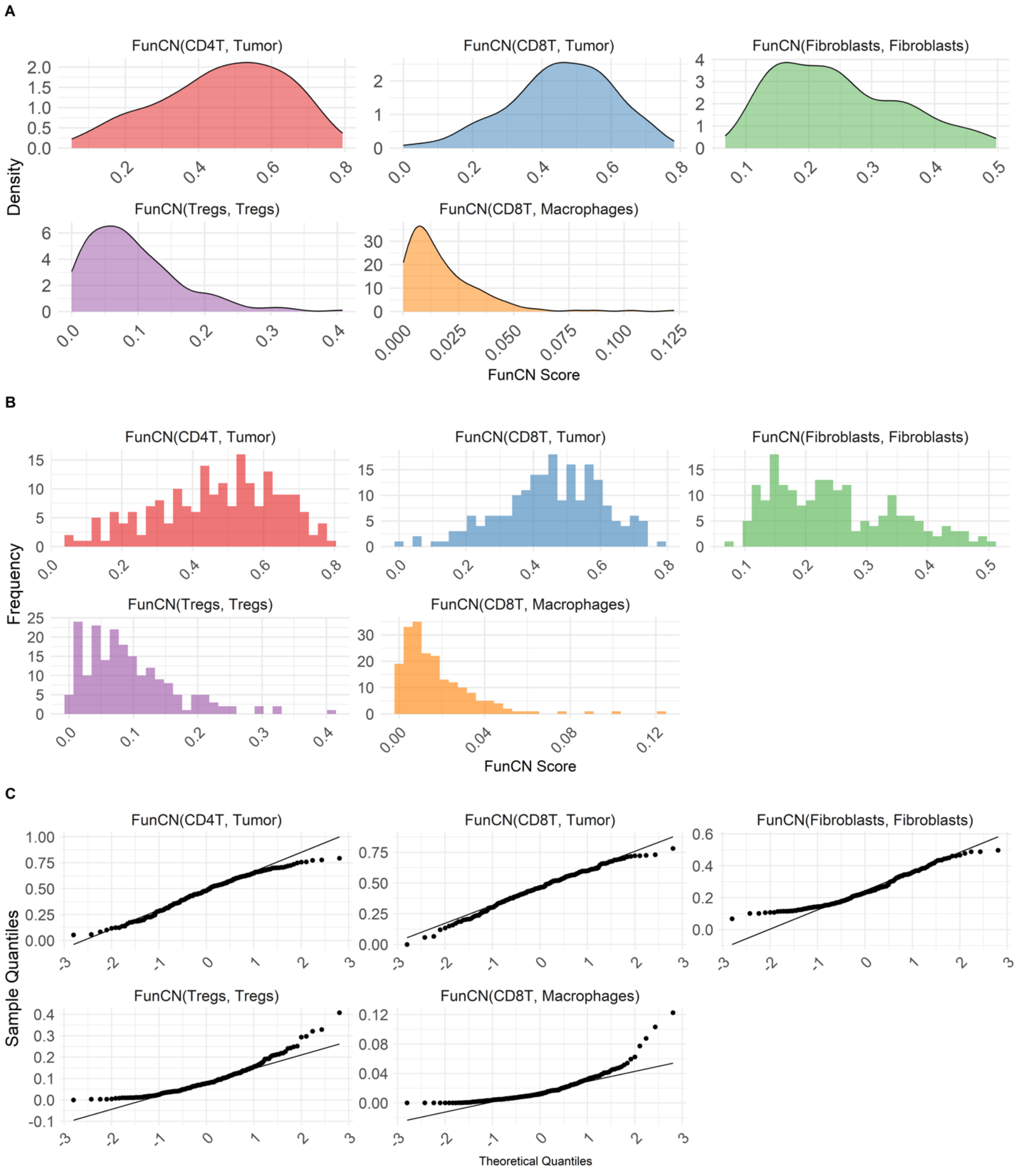


Supplemental Fig. 8: Distribution analysis of five pre-treatment FunCN scores derived from the spQSP model. **(A)** Density plots showing the distribution of FunCN scores for CD4$^{+}$ T cells–tumor, CD8$^{+}$ T cells–tumor, fibroblast–fibroblast, Treg–Treg, and CD8$^{+}$ T cells–macrophage interactions. **(B)** Corresponding frequency histograms for each FunCN score. **(C)** Q–Q plots assessing the normality of each FunCN score distribution by comparing sample quantiles to theoretical normal quantiles.

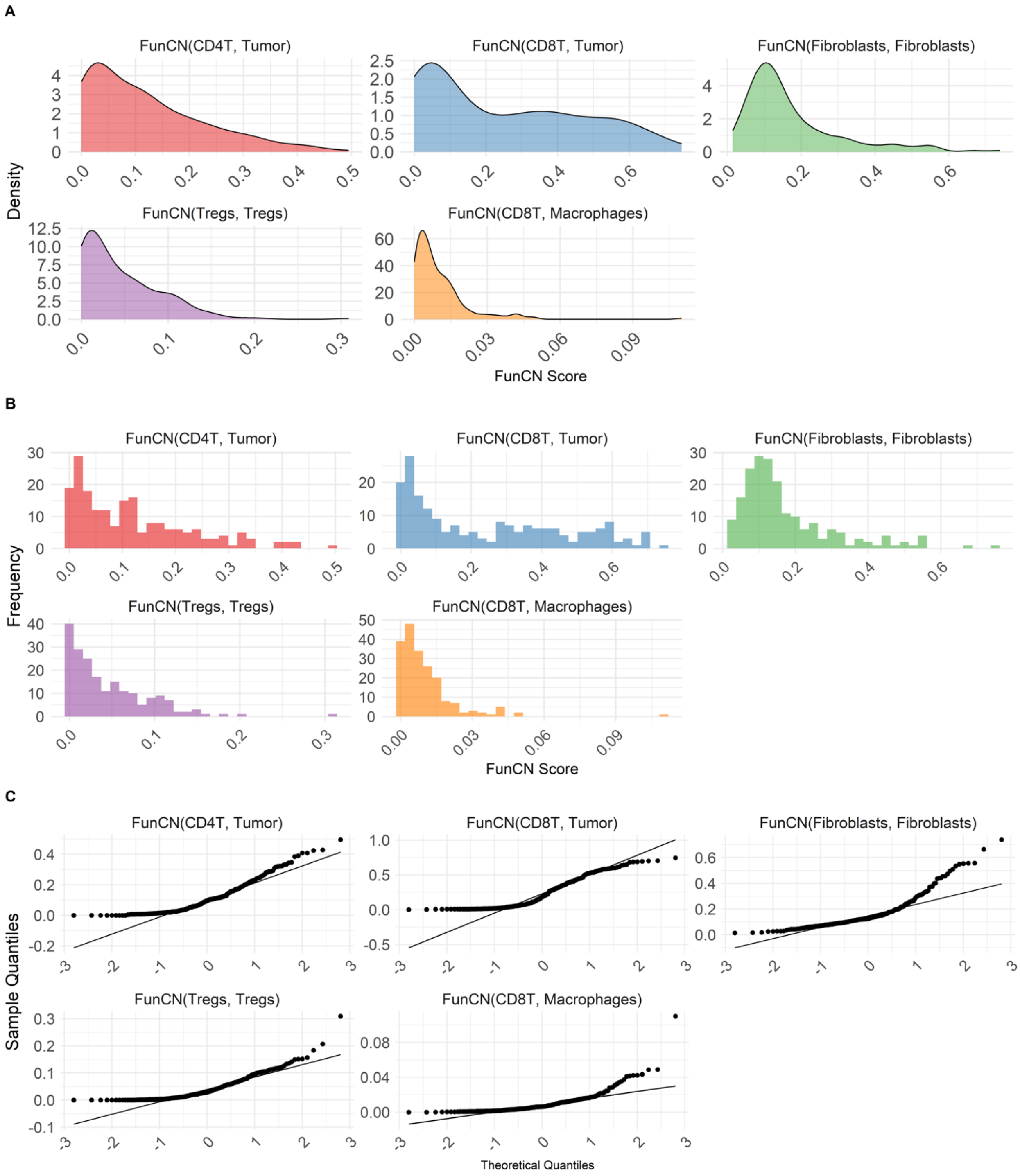


Supplemental Fig. 9: Distribution analysis of five post-treatment FunCN scores simulated by the spQSP model. **(A)** Density plots depicting the distributions of FunCN scores for cell-type pairs: $CD4^+$ T cells–tumor, $CD8^+$ T cells–tumor, fibroblast–fibroblast, Treg–Treg, and $CD8^+$ T cells–macrophages. **(B)** Corresponding histograms showing the frequency distribution of each FunCN score. **(C)** Q–Q plots evaluating the normality of each distribution by comparing sample quantiles to theoretical quantiles.

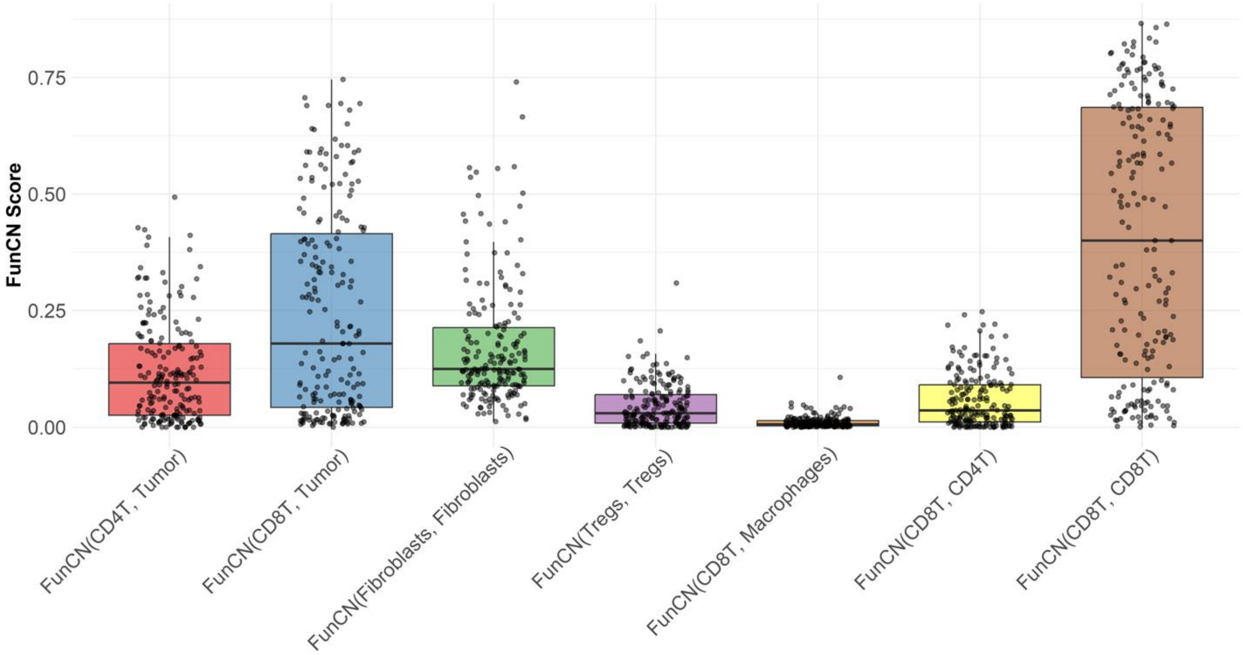


Supplemental Fig. 10: Boxplot of seven post-treatment FunCN scores simulated by the spQSP model. The plot shows the overall distribution of FunCN scores for $CD4^+$ T cells–tumor, $CD8^+$ T cells–tumor, fibroblast–fibroblast, Treg–Treg, $CD8^+$ T cells–macrophage, FunCN CD8T-CD4T, and FunCN CD8T-CD8T interactions. Individual data points are overlaid to visualize sample-level variation within each interaction type.

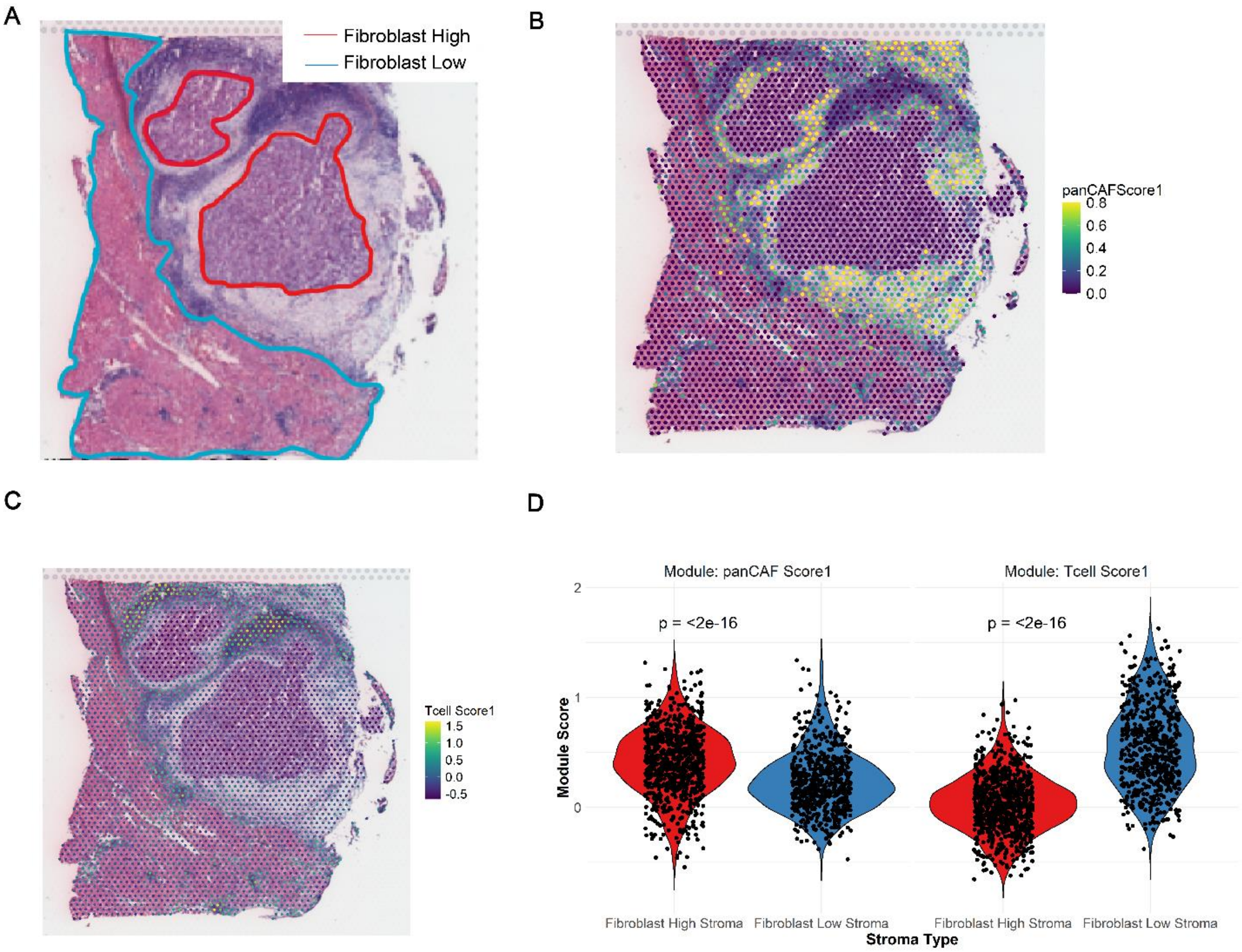


Supplemental Fig. 11: CAF and T cell marker spatial distribution across fibroblast-high and fibroblast-low stroma regions in 10X Visium spatial transcriptomic samples. **(A)** H&E-stained images from Visium slides showing manually outlined tumor regions with high (red) and low (blue) fibroblasts in Sample P8T from Liu et al.. **(B, C)** Spatial distribution of CAF-related module scores (panCAF Score1) and T cell module scores (T cell Score1), overlaid on the corresponding tissue sections in Panel A. CAF module genes: *LUM, DCN, COL1A1, VIM, ENTPD1, ACTA2, PDPN, COL1A2, SERPINF2*; T cell module genes: *CD3D, CD3E, CD8A, CD4, PTPRC, SELL, IL7R , CCL19, CCL21, GZMB.* **(D)** Violin plots showing CAF markers and T cell marker differences in fibroblast high and fibroblast low stroma.

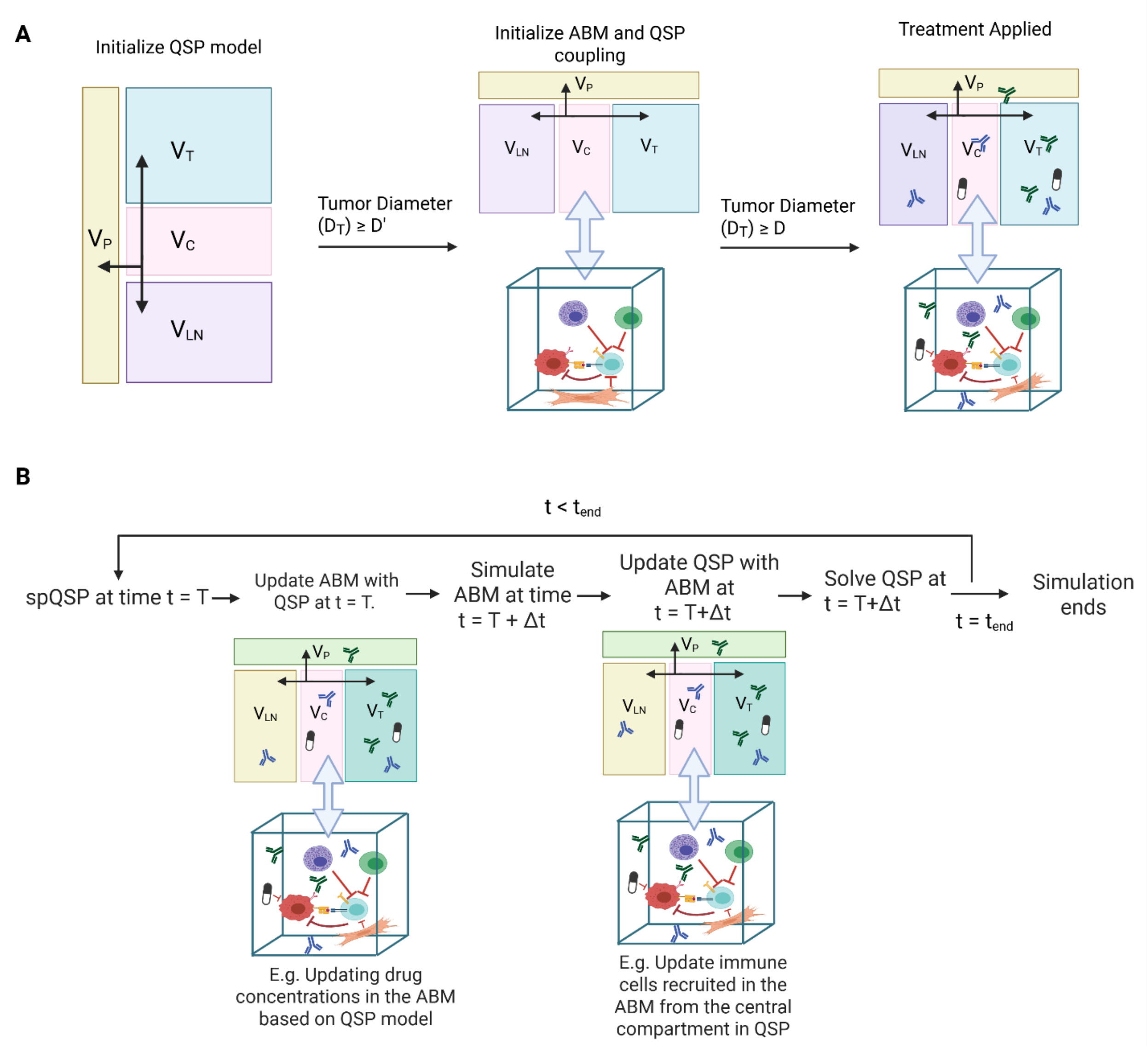


Supplemental Fig. 12 Workflow of the spQSP model. **(A)** Model initialization procedure: The simulation begins with the QSP model alone. When the tumor diameter reaches a threshold $D'$, the agent-based model (ABM) is activated and dynamically coupled with the QSP model. Treatment interventions are applied when the tumor diameter reaches $D$, where $D' = 0.95D$. **(B)** QSP–ABM coupling during simulation: At each simulation timestep, information is exchanged between the QSP model and ABM. Representative examples of this bidirectional coupling are illustrated graphically.
