## Supplemental Methods for "Quantitative Calibration of a Spatial QSP Model Identifies Fibroblast Impact on HCC Immunotherapy"

**Appendix:**

**Agent-Based Model Setup**

The Agent-Based Model (ABM) developed in this study is designed to replicate spatial features observed in multiplexed imaging (e.g., CODEX, IMC) and spatial transcriptomic (ST) datasets. The ABM simulates a 3D tumor microenvironment within a cubic volume of 1 mm × 1 mm × 1 mm, approximating the size of tissue microarrays commonly used for spatial profiling. The simulation domain is discretized into voxels with dimensions of 20 μm × 20 μm × 20 μm. Cells are allowed to migrate to their von Neumann neighborhood (6 adjacent voxels in 3D) via random motility or directed chemotaxis. For cell–cell interactions, cells scan their Moore neighborhood (26 surrounding voxels).

A virtual patient cohort is constructed using Latin Hypercube Sampling (LHS) based on estimated parameter distributions (1). These distributions are calibrated to match cell densities in tumor tissues and clinical overall response rates at the population level (see Supplemental Table 1). Each sampled parameter set defines a unique virtual patient, and each cohort in this study consists of 15 virtual patients. For each virtual patient, the initial tumor diameter $D$ is drawn from a uniform distribution between 3 cm and 7 cm, representing pre-treatment tumor size. The model workflow is summarized in Supplemental Fig. 9A. The simulation begins with a single cancer cell in the QSP model. When the tumor diameter reaches $D^{'}=0.95D$ , the ABM is initialized and dynamically coupled with the QSP model. The two models exchange variables at fixed time intervals ∆t = 6 hours. At each time point T, variables from the QSP model necessary for ABM simulation (e.g., lymphocyte counts in the central compartment) are first passed to the ABM. The ABM is then simulated from $T$ to $T + \Delta t$. Subsequently, output variables from the ABM that inform QSP dynamics (e.g., neoantigen release by tumor cells) are transferred back to the QSP model, which is then simulated over the same interval. This bidirectional coupling ensures model synchronization at each timestep $T + \Delta t$ (see Supplemental Fig. 9B). Therapeutic agents are introduced when the tumor diameter reaches$D$. Simulated spatial outcomes at the end of treatment are compared to spatial proteomic and transcriptomic datasets for validation.

**Tumor Vasculature and Oxygen Delivery**

Our earlier models simulated angiogenesis through the formation of tumor vasculature driven by angiogenic factor gradients(2,3). Various methodologies for modeling tumor vasculature have been comprehensively reviewed(4–6). In the present study, we adopt a simplified yet biologically informed representation of angiogenic factor dynamics, based on prior work (7). Vascular endothelial growth factor (VEGF) is secreted under hypoxic conditions by several tumor-resident cell types. Specifically, progenitor cancer cells secrete VEGF at a rate $k_{sec,VEGF,pro}$ under hypoxic condition ($P_{O_{2},tumor}$< 12 mmHg). M2-like macrophages and stem-like cancer cells also secrete VEGF constantly with rates $k_{sec,VEGF,mac}$ and $k_{sec,VEGF,stem}$, respectively.(8,9) Tumor microvessels uptake VEGF at a rate $k_{uptake,VEGF}.$ The vascular volume fraction $V_{vas}$ represents the volume fraction of tumor microvessels in a given voxel $c\left( x,y,z \right)$, defined as:

$V_{vas}=\frac{\left[ VEGF \right]}{\left[ VEGF \right]+vas_{EC50}}\cdot\left( 1-f_{mooreCancer} \right)\cdot(1-k_{K,cabo})\cdot\frac{[Cabo_{V,T}]}{[Cabo_{V,T}]+IC{50}_{VEGFR2}}$ )	**(1)**

The first term in Eq. 1 is adapted from Mpekris et al (7); the last term reflects pharmacodynamics of cabozantinib(10). $V_{vas}$ is characterized by local concentration of angiogenic factor, $\left[ VEGF \right]$, and fraction of cancer cells in the Moore neighborhood with n=2 (124 closest voxels), $f_{mooreCancer}$. The tumor blood vessels may become non-functional due to compression from surrounding cancer cells(11). $k_{K,cabo}$ is a dimensionless parameter characterizing the anti-angiogenic effect of cabozantinib. Oxygen transport within the tumor is modeled following the Krogh cylinder assumption(12). , where the rate of oxygen delivery is proportional to $V_{vas}$. The temporal evolution of intratumoral oxygen tension is governed by (7,13)

$\frac{\partial P_{O_{2},tumor}}{\partial t}= \nabla\cdot\left( D_{O_{2}}\nabla P_{O_{2},tumor} \right)+{\alpha K}_{v}\cdot\frac{V_{vas}}{A_{vas}}\cdot\left( P_{O_{2}, vas} -P_{O_{2},tumor} \right) -k_{uptake, O_{2}}\cdot C_{voxel}$ **(2)**

$A_{vas}$ is the vascular cross-sectional area per unit volume, with units of $cm^{-1}$; $D_{O_{2}}$is the diffusivity of oxygen, $P_{O_{2}, vas}$is the oxygen partial pressure in the vessels, $k_{uptake, O_{2}}$is the oxygen uptake rate by cancer cell, $C_{voxel}$ is the number of cancer cells in the targeted voxel (which is either 0 or 1). and $K_{v}$ is the is the vascular effective permeability to O_2_ (13)

$K_{v}=2\pi D_{O_{2}}\cdot\frac{\lambda}{\sigma\lambda-\frac{2+\lambda}{4}+\frac{1}{\lambda}ln\frac{1}{w}}$	**(3)**

Here $w=\frac{R_{c}}{R_{t}}$, $\lambda=1-w^{2}$. $R_{c}$ is the vessel radius, and $R_{t}$ is the radius of the Krogh cylinder, estimated as:

$$R_{t}=\frac{1}{\sqrt{\frac{\pi\cdot V_{vas}}{A_{vas}}}}$$

$\sigma=\frac{D_{O_{2}}\alpha_{O2}}{k_{vas}R_{c}}$ is the dimensionless ratio of intravascular to extravascular transport resistance. $k_{vas}$ is the intravascular mass transfer coefficient, and $\alpha_{O2}$ is the oxygen solubility in the tissue.

**Immune Cell Entry Points**

Since the circulating T cells in the blood/central compartment in the QSP model are the source of T cells in the tumor, their recruitment relies on the local vascular density. The probability of a voxel, $c(x,y,z)$ becoming a T cell entry point is

$$p_{entry}=V_{vas}\cdot\frac{\rho_{adhesion}\cdot V_{voxel}}{n_{adhesion}} \mathbf{(4)}$$

Here $\rho_{adhesion}$ is the estimated adhesion molecule density on tumor vasculature; $V_{voxel}$ is volume per voxel, and $n_{adhesion}$ is the number of adhesion molecules required for recruiting an immune cell. The immune cell entry points are calculated iteratively based on Eq. 4 at the beginning of each time step.

**Macrophage Recruitment**

We extended the macrophage module from Wang et al.(14) to make a spatial version, which in turn is a simplified mechanistic model of macrophage polarization developed by Zhao et al.(15). The Arg1+ macrophage module is modified from our previous models(16,17). Arg1 negative macrophage has two states: M1 and M2 (referred as macrophage). M1 macrophages secrete pro-inflammatory cytokines, including IFNγ and IL12, whereas M2 macrophages release immunosuppressive cytokines (IL10 and TGFβ) and angiogenic factor VEGFA. Arg1+ macrophage releases anti-inflammatory cytokines including Arginase-I (ArgI) and immunosuppressive molecule nitric oxide (NO). In the QSP model, the recruitment rate of macrophages into the tumor is:

$$\frac{dM}{dt}=k_{Mac,recr}\cdot V_{T}\cdot\frac{\left[ CCL2 \right]}{\left[ CCL2 \right]+CCL2_{EC50}} \mathbf{(5)}$$

Here**,** $M$ is the total number of macrophages, $k_{Mac,recr}$ is a parameter abstracting both monocyte extravasation rate and transformation rate to M1-like macrophage; $V_{T}$ is the volume of the tumor, and $CCL2_{EC50}$ is the half-maximal CCL2 macrophage recruitment. To translate the recruitment mechanism into the ABM module, the probability of recruiting a macrophage at every voxel over the time step interval $\Delta t$ is:

$$p_{Mac, rec}=k_{Mac,recr}\cdot V_{voxel}\cdot\frac{\left[ CCL2 \right]}{\left[ CCL2 \right]+CCL2_{EC50}}\cdot\Delta t \mathbf{(6)}$$

$$p_{MDSC, rec}=k_{MDSC,recr}\cdot V_{voxel}\cdot\frac{\left[ CCL2 \right]}{\left[ CCL2 \right]+CCL2_{EC50}}\cdot\Delta t \mathbf{(7)}$$

**Macrophage Death**
In the QSP model, macrophage death (for both M1 and M2 macrophages) is defined by the following equation:

$$\frac{dMac}{dt}=-k_{Mac,death}Mac \mathbf{(8)}$$

where $k_{Mac,death}$ is the death rate of macrophages, we models the survival time distribution of recruited macrophage as an exponential distribution

$$T_{mac} \sim Exp(\frac{1}{k_{Mac,death}}) \mathbf{(9)}$$

**M1 to M2 Macrophage Polarization**

The QSP model formulates M1 to M2 polarization as:

$\frac{dM_{1}}{dt}= -k_{pol, M1}\cdot\frac{\left[ TGFB \right]}{\left[ TGFB \right]+TGFB_{EC50}}\cdot\frac{\left[ IL10 \right]}{\left[ IL10 \right]+{IL10}_{EC50}}\cdot M_{1}$ $\mathbf{(10)}$

which translates to $p_{M1 to M2 polarization}=1-e^{-\alpha_{M1,polar}\Delta t}$for individual M1 macrophage polarized to M2 macrophage. $\alpha_{M1,polar}$ denotes the propensity of M1-like macrophage polarizing into M2-like macrophage:

$$\alpha_{M1,polar}= k_{pol, M1}\cdot\frac{[TGFB]}{[TGFB]+TGFB_{EC50}}\cdot\frac{[IL10]}{[IL10]+{IL10}_{EC50}} \mathbf{(11)}$$

**M2 to M1 Macrophage Polarization**

The QSP model formulates M2 to M1 polarization as:

$$\frac{dM_{2}}{dt}= -k_{pol, M2}\cdot\frac{\left[ IL12 \right]}{\left[ IL12 \right]+{IL12}_{EC50}}\cdot\frac{\left[ IFN\gamma\right]}{\left[ IFN\gamma\right]+{IFN\gamma}_{EC50}}\cdot M_{2} \mathbf{(12)}$$

which translates to $p_{M2 to M1 polarization}=1-e^{-\alpha_{M2,polar}\tau}$for individual M1 macrophage polarized to M2 macrophage$. \alpha_{M2,polar}$ denotes the propensity of M2-like macrophage polarizing into M1-like macrophage:

$$\alpha_{M2,polar}= k_{pol, M2}\cdot\frac{[IL12]}{[IL12]+{IL12}_{EC50}}\cdot\frac{[IFN\gamma]}{[IFN\gamma]+{IFN\gamma}_{EC50, M1}} \mathbf{(13)}$$

**M1 Macrophage Phagocytosis**

M1 macrophage mediated phagocytosis of cancer cell is modeled in the QSP model as:

$$\frac{dC}{dt}= -k_{phago}\cdot\frac{M_{1}}{M_{1}+C+1}\cdot(1-H_{Mac, C1})\cdot(1-H_{IL10,phago}) \mathbf{(14)}$$

The phagocytosis mechanism in ABM model is translated to $p_{phago}=1-e^{-\alpha\Delta t}$ , where $\alpha_{phago}$ is defined as the propensity of a cancer cell phagocytosed by nearby M1-like macrophage:

$$\alpha_{phago}=k_{phago}\cdot\frac{N_{M_{1}}}{N_{M_{1}}+N_{C}+1}\cdot\left( 1-H_{Mac, C1} \right)\cdot\left( 1-H_{IL10,phago} \right) \mathbf{(15)}$$

$N_{M_{1}}$ and $N_{C}$ are number of adjacent M1-like macrophages and cancer cells to the target cancer cell, respectively. $k_{phago}$ represents M1-like macrophage phagocytosis rate; $H_{IL10,phago}$ is the Hill function for IL10 inhibition of phagocytosis, and $IL{10}_{phago, EC50}$ is the half-maximal IL-10 level for inhibition of phagocytosis by macrophage.

$$H_{IL10}=\frac{\left[ IL10 \right]^{2}}{\left[ IL10 \right]^{2}+{IL10}_{phago, EC50}^{2}}$$

$H_{Mac, C1}$ is the total Hill function for checkpoint inhibition of phagocytosis, which constitutes $SIRP\alpha$ mediated and PD1 mediated inhibition:

$$H_{Mac, C}= 1-\left( 1-H_{SIRP\alpha} \right)\cdot\left( 1-H_{PD1} \right) \mathbf{(16)}$$

The PD1 induced checkpoint inhibition ($H_{PD1}$) is identical to the inhibition mechanism in the T cell module, and $H_{SIRP\alpha}$ is the Hill function of SIRPα induced inhibition, written as:

$$H_{SIRP\alpha}=\frac{\left[ CD47SIRP\alpha\right]^{2}}{\left[ CD47SIRP\alpha\right]^{2}+SIRP\alpha_{EC50}^{2}} \mathbf{(17)}$$

The binding dynamic of CD47 and $SIRP\alpha$ is described as:

$$k_{on, SIRP\alpha}\left[ CD47 \right]\left[ SIRP\alpha\right]\underset{\leftrightarrow}{}k_{off, SIRP\alpha}\left[ CD47SIRP\alpha\right] \mathbf{(18)}$$

At steady state:

$$k_{on, SIRP\alpha}\left( \left[ CD{47}_{Cancer} \right]-\left[ CD47SIRP\alpha\right] \right)\cdot\left( \left[ SIRP\alpha_{Mac} \right]-\left[ CD47SIRP\alpha\right] \right)=k_{off, SIRP\alpha}\left[ CD47SIRP\alpha\right] \mathbf{(19)}$$

$$\frac{k_{on, SIRP\alpha}}{k_{off, SIRP\alpha}}{(\left[ CD47SIRP\alpha\right]}^{2}-(\left[ CD47 \right]+ \left[ SIRP\alpha\right]+ \frac{k_{off, SIRP\alpha}}{k_{on, SIRP\alpha}})\cdot\left[ CD47SIRP\alpha\right]+\left[ CD47 \right]\left[ SIRP\alpha\right])=0 \mathbf{(20)}$$

If we denote $\left[ CD47 \right]=a; \left[ SIRP\alpha\right]=b; \left[ CD47SIRP\alpha\right]=x$, then

$$x^{2}-\left( a+b+\frac{k_{off, SIRP\alpha}}{k_{on, SIRP\alpha}} \right)x+ab=0 \mathbf{(21)}$$

Based on values of a, b, $k_{off, SIRP\alpha}$, and $k_{on, SIRP\alpha}$, the equation has a real root that is larger than zero and less than both $a$and b. The root is then used for calculating $H_{SIRP\alpha}$ in Eq. 17.

**Cancer Cell Proliferation**

The QSP module characterizes the cancer cell proliferation as a logistic growth, written as $C$ is the cancer cell counts in the QSP model:

${\frac{dC}{dt}=k}_{C, growth}\cdot(1-k_{K,cabo}\cdot\frac{[cabo_{T}]}{[cabo_{T}]+IC{50}_{MET}})\cdot(C_{max}-C)\cdot C$ $\mathbf{(22)}$

Therefore, the division time (doubling time) of cancer cell is calculated as:

$$t_{cancer, division}=\frac{\ln\left( 2 \right)}{k_{C, growth}\cdot(1-k_{K,cabo}\cdot\frac{[cabo_{T}]}{[cabo_{T}]+IC{50}_{MET}})} \mathbf{(23)}$$

Here $\lambda_{c}$ is a positive parameter modeling the effect of cabozantinib inhibiting MET pathway which reduces cancer proliferation rate. If cancer cells are in hypoxic condition ($P_{O_{2},tumor}$< 12 mmHg), $t_{cancer, division}$ is doubled.

**Coupling of QSP and ABM Modules**
The spQSP model simulates dynamical changes of lymph nodes and central compartment in the QSP module, and the ABM module is constructed specifically to represent a spatially resolved volume in the tumor compartment. As the ABM module simulates anti-tumoral immunity dynamics of the region of interest (ROI) in the tumor microenvironment, the rest part of the tumor is delegated to the QSP module. We assign a weight, $w_{QSP}$, representing portion of the tumor simulated in the QSP model, and the remaining ${1-w}_{QSP}$ is represented by the ABM module. Also, information exchange between A is reflected in the ABM module. For instance, recruitment of CD8+ T cells and Treg from the central compartment and antigen transportation from the tumor to the lymph node requires coupling between the ABM and the QSP module. The scaling factor *s* of the ROI is calculated based on following equation:

$$s=\frac{1-w_{QSP}}{w_{QSP}}\cdot\frac{C_{QSP}}{C_{ABM}} \mathbf{(1)}$$

where *C_QSP_* and *C_ABM_* are the number of cancer cells in the QSP module and ABM module, respectively. We track the number of recruited immune cells and the amount of tumor neoantigen produced in the ABM module during simulation. These quantities are then multiplied by the scaling factor *s* and updated in the QSP module at each timestep.

**Molecular Agent Transport in ABM module**

The ABM module is composed of both cellular components and molecular components. Molecular components, including cytokine and nutrients, are secreted by cell agents. The partial differential equation (PDE) governing the concentration of each molecular agent $m$ is defined as:

$$\frac{\partial m}{\partial t}=D\nabla^{2}m-\mu m+S\left( x,y,z,t \right)-U\left( x,y,z,t \right) \mathbf{(2)}$$

where *D* is molecular diffusivity, *μ* is the degradation rate, and $S$ is the secretion rate of $m$ at voxel (*x, y, z*) with time *t*, and $U$ is the consumption rate of $m$ at voxel (*x, y, z*) with time *t*.

We use finite volume method with no flux boundary condition implemented in the BioFVM software to solve PDE equations(18). Spatial discretization of the molecular component matches the voxels in the agent layer. When molecular agent $m$ is released by a cell, a source of the factor is created at the location corresponding to the center of the cell’s voxel. Conversely, a sink is created as $m$ is consumed by a cell.

**Discretization of Continuous ODE equations:**

Many cell behaviors in the QSP model, involving cell death and cell state transformation are modeled by an exponential decay function, written as:

$$S=S_{0}\cdot exp\left( -k_{d}\tau\right) \mathbf{(3)}$$

$S_{0}$ denotes cell amount at initial time $t_{0}$; $k_{d}$ is a decay constant; $S$ is the cell amount at time $t_{0}+\tau$. An intermediate step representing fraction of decayed cell for a period of time $\tau$ is:

$$\frac{S_{0}-S}{S_{0}}=1-exp\left( -k_{d}\tau\right) \mathbf{(4)}$$

Therefore, to reflect decay behavior in the ABM model, the degradation probability of a cell is:

$$p_{decay}=1-e^{-\alpha\tau}$$

Where $\alpha$ can be either a constant or a function charactering more complex behaviors.

**Cancer Cell Initiation:**

Cancer cells have only one state in the QSP model. The cancer progression is governed by the growth rate, natural death rate, and immune cell killing rate. In ABM module, the cancer cells are differentiated into cancer stem-like cells, progenitor cells, and senescent cells, and rules are adapted from our previous models(19,20). Stem-like cancer cells have unlimited number of division cycles with a rate *r_s_*, with asymmetric division rate, *k*. The asymmetric division produce one daughter progenitor cell and one daughter stem-like cancer cell. The probability of symmetric division is *1-k*, which generate two daughter stem-like cancer cells. Progenitor cancer cells, with limited number of divisions ($d_{max}$), divide at a rate *r_p_*. The parent cells become senescent after $d_{max}$ rounds of division. Senescent cells cannot proliferate and have death rate of *µ*. The reproduction rates of both stem-like cancer cells and death rate of senescent cells are derived from QSP model parameters. *S_c_*, *P_i_* and *S_n_* denote cancer stem-like cancer cell, progenitor cell after *i* divisions, and senescent cell, respectively.

$$\frac{dS_{c}}{dt}= r_{s}\left( 1-k \right)S_{c} \mathbf{(5)}$$

$$\frac{dP_{1}}{dt}= r_{s}kS_{c}- r_{p}P_{1} \mathbf{(6)}$$

$$\frac{dP_{i}}{dt}=2r_{p}P_{i-1}-r_{p}P_{i} for 2\leq i\leq dmax \mathbf{(7)}$$

$$\frac{dS_{n}}{dt}=2r_{p}P_{dmax}-\mu S_{n} \mathbf{(8)}$$

Combining equations (**5**) and (**6**):

$\frac{d}{dt}\left( P_{1}-pS_{c} \right)=-r_{p}(P_{1}-pS_{c})$	$\mathbf{(9)}$

Where,

$$p=\frac{kr_{s}}{r_{s}\left( 1-k \right)+r_{p}}$$

$\left( pS_{c}-P_{1} \right)$ approaches 0, as cancer cell counts approaches infinity. The system asymptotically approaches a stable state where all cancer species grow with same rate *r**. The ratio among cancer cell species asymptomatically approaches the constant *r**:

$$r^{*}=r_{s}\left( 1-k \right)$$

$$\frac{P_{1}\left( t \right)}{S_{c}\left( t \right)}=\frac{kr_{s}}{r^{*}+r_{p}}=l_{0} \mathbf{(10)}$$

$$\frac{P_{i}\left( t \right)}{P_{i-1}\left( t \right)}=\frac{2r_{p}}{r^{*}+r_{p}}=l_{1}, 1<i\leq dmax \mathbf{(11)}$$

$$\frac{S_{n}\left( t \right)}{P_{dmax}\left( t \right)}=\frac{2r_{p}}{r^{*}+\mu}=l_{2} \mathbf{(12)}$$

As all cancer cell species’ growth rates approach constant *r**, the growth rate equals cancer cell growth rate from QSP module, defined in the QSP parameter file. *r** can be expressed as a function of *k* and $r_{s}$, when *k* is a constant, the growth rate of CSC becomes$r_{s}= \frac{r^{*}}{1-k}$ and $r_{p}$ and *µ* can be set independently. Based on results from equations (**10**) - (**12**), at steady state, the ratio between any pair of cancer cell species remains as constant while tumor is expanding. The number of cancer cells initialized in the ABM module is linearly proportional to $D_{init}^{3}$, where $D_{init}$ is the initial diameter of tumor. The fraction of each cancer cell species when ABM module is initialized is:

| Species | Fraction |
| --- | --- |
| $S_{c}$ | $\frac{1}{q}$ |
| $P_{i}$ | $\frac{pl_{1}^{i-1}}{q}$ |
| $S_{n}$ | $\frac{pl_{1}^{dmax-1}l_{2}}{q}$ |

where$q=1+p\frac{l_{1}^{dmax}-1}{l_{1}-1}+pl_{1}^{dmax-1}l_{2}$

**CD8 T Cell Recruitment:**

For each CD8 T cell entry point, the probability of recruiting a CD8 T cell into the ABM model is written as

$$p_{CD8T,rec}={T_{CD8T,central}\cdot k}_{CD8T, rec}\cdot V_{voxel}\cdot\tau$$

Where $T_{CD8T,central}$is CD8+ T cell count in the central compartment $k_{T, rec}$ is the T cell recruitment rate into the tumor. $V_{voxel}$ is the volume of each single voxel, and $\tau$ is the time per time step.

**CD8+ T cell killing of cancer cells**
In the QSP module, the CD8+ T cell killing rate is defined as:

$\frac{dC}{dt}=- k_{CD8,killing} \cdot\frac{T_{CD8}}{C +Tcell+cell}\left( 1-H_{PD1} \right)\left( 1-H_{MDSC,c} \right)C \mathbf{(15)}$

Where $k_{CD8,killing}$ is the cancer cell killing rate by T cell. To translate that equation to the ABM module, the probability of a cancer cell killing by adjacent CD8+ T cell is calculated as:

$$p_{kill}= 1-e^{-\Delta t\cdot\alpha_{c,CD8}}$$

where $\Delta t$ is the length of each time step, and $\alpha_{c, CD8}$ is defined as:

$$\alpha_{c, CD8}=k_{CD8,killing} \cdot\frac{N_{CD8}}{N_{c} + N_{CD8}+N_{cell}}\left( 1-H_{PD1} \right)\left( 1-H_{MDSC,c} \right)$$

$\frac{N_{CD8}}{N_{c} + N_{CD8}+N_{cell}}$ is the fraction of cytotoxic T cells among all cells in the Moore neighborhood of the target cancer cell.

**CD8+ T cell exhaustion**
CD8+ T cells are exhausted by two mechanism in the spQSP model: PDL1 mediated exhaustion and Treg mediated exhaustion.
In the QSP module, the rate of CD8+ T cell being exhausted by PD1 - PDL1 interaction is:

$$\frac{dT_{CD8}}{dt}=- T_{CD8}* k_{ex,PDL1}\cdot\frac{C}{C +Tcell+cell}\cdot H_{PD1}\left( X, 1 \right) \mathbf{(16)}$$

where $cell$ is the counts of cells other than T cell or cancer cell, and $\frac{C}{C +Tcell+cell}$ is the fraction of cancer cells in the tumor. In the ABM module the probability of single CD8+ T cell being exhausted by PD1 - PD-L1 interaction is:

$p_{exhaust, PDL1}=1-e^{-\Delta t \cdot k_{ex,PDL1} \cdot H_{PD1}(X,1))*q_{c}} =1-{B_{PDL1}}^{H_{PD1}\left( X,1 \right)q_{c}}$ $\mathbf{(17)}$
here, $B_{PDL1}=e^{-\Delta t \cdot k_{ex,PDL1}}$, and $q_{c}$ is the fraction of cancer cells among all cells in the Moore neighborhood of the target CD8+ T cell. Since we assume all cancer cells express PD-L1 in the ABM module, $q_{c}$ equals to 1.

To account the rate of CD8+ T cell being exhausted by Treg is:
 $\frac{dT_{CD8}}{dt}=- T_{CD8}\cdot k_{ex,Treg}\cdot\frac{Treg}{C +Tcell+cell}$ $\mathbf{(18)}$In the ABM module the probability of single CD8+ T cell being exhausted by Treg is:

$$p_{exhaust, Treg}=1-e^{-\Delta t\cdot k_{ex,Treg}\cdot q_{Treg}} \mathbf{(19)}$$

Here $q_{Treg}$is the fraction of Tregs in the Moore neighborhood of targeted CD8+ T cell.

**CD4 T cell recruitment:**

The entry point mechanism for CD4 T cell, including both regulatory T cell (Treg) and helper T cell (Th) is the same. For each CD4 T cell entry point, the probability of either Th or Treg cell into the ABM model is written as

$$p_{Treg,rec}={T_{reg,central}\cdot k}_{CD4, rec}\cdot V_{voxel}\cdot\tau$$

$$p_{Th,rec}={T_{h,central}\cdot k}_{CD4, rec}\cdot V_{voxel}\cdot\tau$$

Where $T_{Treg,central}$and $T_{Th,central}$ are Treg and Th counts in the central compartment, respectively. $k_{CD4, rec}$ is the CD4 T cell recruitment rate into the tumor.

**CD4 T cell expansion and death:**

In the ABM module, the expansion of regulatory T cell induced by $ArgI$ at each timestep is defined:

$$p_{exp, Treg, ArgI}=\frac{[ArgI]}{[ArgI]+{EC50}_{ArgI,Treg}} \mathbf{(20)}$$

The death rate of CD4 T cell in the QSP model is defined as:

$${{\frac{dT_{CD4}}{dt}=-k}_{TCD4,death}\cdot T}_{CD4}$$

Where $k_{TCD4,death}$ is the death rate of CD4 T cell. Therefore, we model the survival time distribution of recruited CD4 T cell as an exponential distribution

$$T_{TCD4} \sim Exp(\frac{1}{k_{TCD4,death}})$$

**Modeling of PD-1–PDL1 interaction**

We assume all CD8+ T cell express PD-1 and all cancer cells express PDL1; this assumption can be readily changed to fit experimental data where available. Our model represents PDL1 and PD1 interactions in multiple ways. The model takes the number of PD1—PDL1 bonds in the immune synapse as input to the following Hill function:

$$H_{PD1}\left( X,n \right)=\frac{{(\frac{X}{k_{PDL1,PD1}})}^{n}}{1+{(\frac{X}{k_{PDL1,PD1}})}^{n}} \mathbf{(21)}$$

where $X$ is the total number of PD-1 – PDL1 bonds between CD8+ T cell and cancer cell.
The number of PD-1 molecules in the immune synapse can be calculated as:

$$[PD1_{T}] =N_{PD1, T} \cdot\frac{A_{syn}}{A_{T}} \mathbf{(22)}$$

Total number of PD-L1 molecules involved in the immune synapse can be calculated as:

$$N_{PDL1,max} = N_{PDL1,cancer}\cdot\frac{A_{syn}}{A_{c}}$$

$$\left[ PDL1_{cancer} \right]= N_{PDL1,max}\cdot\frac{\left[ IFN\gamma\right]}{\left[ IFN\gamma\right]+{IFN\gamma}_{EC50, PDL1}} \mathbf{(23)}$$

Where ${IFN\gamma}_{EC50, PDL1}$ is the half-maximal $IFN\gamma$ concentration required for PDL1 expression on cancer cell. Assuming immune synapse reaches equilibrium:

$$[PDL1PD1]= [PD1]\cdot[PDL1_{cancer}]\cdot k_{1}$$

$$\left[ PD1Nivo \right]= [PD1]*[Nivo]\cdot k_{2}$$

$$\left[ PD1NivoPD1 \right]= \left[ PD1Nivo \right]\cdot[PD1]\cdot k_{3}$$

$$[PDL1PD1]+[PD1]+\left[ PD1Nivo \right]+2\cdot\left[ PD1NivoPD1 \right]= [PD1_{T}] =T_{1}$$

$$[PDL1PD1]+[PDL1]= [PDL1_{cancer}] =T_{2}$$

where

$$k_{1}= \frac{k_{on, PD1,PDL1}}{k_{off, PD1,PDL1} *A_{syn}}$$

$$k_{2}= \frac{2*k_{on,Nivo}}{k_{off,Nivo}*\gamma_{T, Nivo}}$$

$$k_{3}=\frac{X_{PD1, Nivo}*k_{on, PD1Nivo}}{2*k_{off, PD1Nivo}}$$

Here $X_{PD1, Nivo}$ is the antibody cross-arm binding efficiency, and $\gamma_{T, Nivo}$ is the volume fraction available to nivolumab.

Let $[PDL1PD1]$ = *X* and x = *X/T_2._* We rewrite the equation as:

$x^{3}-\left( 2+\frac{1}{k_{1}T_{2}}+\frac{T_{1}}{T_{2}}+\frac{[Nivo]*k_{2}}{k_{1}T_{2}}\left( 1-\frac{2k_{3}}{k_{1}} \right) \right)x^{2}+\left( 1+\frac{1}{k_{1}T_{2}}+\frac{{2T}_{1}}{T_{2}}+\frac{[Nivo]*k_{2}}{k_{1}T_{2}} \right)x-\frac{T_{1}}{T_{2}}=0$ $\mathbf{(24)}$

During simulation, equation **(24)** is solved respect to $x$ using Newton–Raphson method with initial guessing point $x_{0}=0$ at every timestep $t$. Then $[PDL1PD1]$, which equals to *xT2*, during immune synapse is calculated dynamically during simulation when each CD8+ T cell interacts with cancer cells. It can be shown that for the parameters of the problem the roots of the equation are real, and we choose the root in the interval 0<x<1.

**Reference:**

1.	Cheng Y, Straube R, Alnaif AE, Huang L, Leil TA, Schmidt BJ. Virtual Populations for Quantitative Systems Pharmacology Models. Methods Mol Biol. 2022;2486:129–79.

2.	Norton K-AA, Jin K, Popel AS. Modeling triple-negative breast cancer heterogeneity: Effects of stromal macrophages, fibroblasts and tumor vasculature. J Theor Biol. 2018 Sep;452:56–68.

3.	Norton KA, Popel AS. Effects of endothelial cell proliferation and migration rates in a computational model of sprouting angiogenesis. Sci Rep. 2016;6(October):1–10.

4.	Apeldoorn C, Safaei S, Paton J, Maso Talou GD. Computational models for generating microvascular structures: Investigations beyond medical imaging resolution. WIREs Mech Dis. 2023;15(1):1–17.

5.	Zhang Y, Wang H, Oliveira RHM, Zhao C, Popel AS. Systems biology of angiogenesis signaling: Computational models and omics. WIREs Mech Dis. 2021;(December):1–37.

6.	Hormuth DA, Phillips CM, Wu C, Lima EABF, Lorenzo G, Jha PK, et al. Biologically-based mathematical modeling of tumor vasculature and angiogenesis via time-resolved imaging data. Cancers (Basel). 2021;13(12).

7.	Mpekris F, Voutouri C, Baish JW, Duda DG, Munn LL, Stylianopoulos T, et al. Combining microenvironment normalization strategies to improve cancer immunotherapy. Proc Natl Acad Sci U S A. 2020;117(7):3728–37.

8.	Afify SM, Sanchez Calle A, Hassan G, Kumon K, Nawara HM, Zahra MH, et al. A novel model of liver cancer stem cells developed from induced pluripotent stem cells. Br J Cancer. 2020;122(9):1378–90.

9.	Wu WK, Llewellyn OPC, Bates DO, Nicholson LB, Dick AD. IL-10 regulation of macrophage VEGF production is dependent on macrophage polarisation and hypoxia. Immunobiology. 2010;215(9–10):796–803.

10.	Hutchinson LG, Mueller HJ, Gaffney EA, Maini PK, Wagg J, Phipps A, et al. Modeling longitudinal preclinical tumor size data to identify transient dynamics in tumor response to Antiangiogenic Drugs. CPT Pharmacometrics Syst Pharmacol. 2016;5(11):636–45.

11.	Griffon-Etienne G, Boucher Y, Brekken C, Suit HD, Jain RK. Taxane-induced apoptosis decompresses blood vessels and lowers interstitial fluid pressure in solid tumors: clinical implications. Cancer Res. 1999 Aug;59(15):3776–82.

12.	Sharan M, Gupta S, Popel AS. Parametric analysis of the relationship between end-capillary and mean tissue PO2 as predicted by a mathematical model. J Theor Biol. 1998;195(4):439–49.

13.	Popel AS. Theory of oxygen transport to tissue. Crit Rev Biomed Eng. 1989;17(3):257–321.

14.	Wang H, Zhao C, Santa-Maria CA, Emens LA, Popel AS. Dynamics of tumor-associated macrophages in a quantitative systems pharmacology model of immunotherapy in triple-negative breast cancer. iScience. 2022;25(8):104702.

15.	Zhao C, Medeiros TX, Sové RJ, Annex BH, Popel AS. A data-driven computational model enables integrative and mechanistic characterization of dynamic macrophage polarization. iScience. 2021;102112.

16.	Wang H, Sové RJ, Jafarnejad M, Rahmeh S, Jaffee EM, Stearns V, et al. Conducting a Virtual Clinical Trial in HER2-Negative Breast Cancer Using a Quantitative Systems Pharmacology Model With an Epigenetic Modulator and Immune Checkpoint Inhibitors. Front Bioeng Biotechnol. 2020;8(February):1–16.

17.	Ruiz-Martinez A, Gong C, Wang H, Sove RJ, Mi H, Kimko H, et al. Simulations of tumor growth and response to immunotherapy by coupling a spatial agent-based model with a whole-patient quantitative systems pharmacology model. PLoS Comput Biol. 2022 Jul;18(7):1–32.

18.	Ghaffarizadeh A, Friedman SH, MacKlin P. BioFVM: An efficient, parallelized diffusive transport solver for 3-D biological simulations. Bioinformatics. 2016;32(8):1256–8.

19.	Norton KA, Jin K, Popel AS. Modeling triple-negative breast cancer heterogeneity: Effects of stromal macrophages, fibroblasts and tumor vasculature. J Theor Biol. 2018;452:56–68.

20.	Gong C, Milberg O, Wang B, Vicini P, Narwal R, Roskos L, et al. A computational multiscale agent-based model for simulating spatio-temporal tumour immune response to PD1 and PDL1 inhibition. J R Soc Interface. 2017;14(134).
